# Hemin-driven chromatin remodelling by atherosclerotic risk gene *SMARCA4* switches human blood-derived macrophages from leukocyte disposal to erythrocyte disposal

**DOI:** 10.1101/2023.05.01.538808

**Authors:** Luke Cave, Katharine M Lodge, Derick Chiappo, Shivani Sinha, Faiz Chughtai, Adam Tsao, Dorian O Haskard, Justin C Mason, Steve E Humphries, Joseph J Boyle

**Affiliations:** National Heart and Lung Institute (NHLI), Imperial College London; Centre for Cardiovascular Genetics, Institute for Cardiovascular Science, University College London, London

## Abstract

**Background:** Putative genetic risk loci for atherosclerotic vascular disease include *SMARCA4*, a chromatin remodeling gene important for gene activation. Its causal role in atherosclerosis has been uncertain. Intraplaque hemorrhage (IPH) is a late event in atherosclerosis that is linked to plaque destabilisation and increased inflammation. IPH is countered by Mhem macrophages, which are directed by hemin-mediated induction of Heme Oxygenase 1 (*HMOX1*) via Activating Transcription Factor 1 (ATF1). *Atf1* deficiency *in vivo* impairs hematoma clearance, promoting inflammation and oxidative stress. Like its homologue cyclic-adenosine monophosphate response element binding protein 1 (CREB1), ATF1 is normally cyclic-AMP activated.

**Hypothesis:** Hemin-directed chromatin remodelling by SMARCA4 regulates specificity of ATF1 gene-binding, thereby switching between leukocyte disposal and erythrocyte disposal, contributing to its role in atherosclerosis.

**Results:** We here show that *SMARCA4* is genetically independent of the adjacent *LDLR* locus (p<0.05). In human blood-derived macrophages, hemin triggered histone acetylation (H3K9Ac) and SMARCA4 recruitment in advance of p-ATF1 recruitment at the *HMOX1* enhancer. si-RNA-mediated *SMARCA4*-knockdown suppressed p-ATF1 binding to *HMOX1* but increased its binding to cyclic-AMP responsive genes *FOS* and *NR4A2*, with corresponding changes in mRNA levels. This functionally correlated with *SMARCA4*-knockdown switching hemin to mimic prostacyclin (PGI_2_), for induced genes and phagocytic disposal of leukocytes rather than erythrocytes.

**Conclusions:** These data establish *SMARCA4* as an independent atherosclerosis risk gene and reveal a novel mechanism in which it switches between disposal of leukocytes or erythrocytes, with important clinical implications for atherosclerotic inflammation and intraplaque hemorrhage including treatment by histone deacetylase inhibitors.

## Introduction

Atherosclerosis is a vascular pathology that causes enormous levels of morbidity and mortality including myocardial infarction and stroke^1^. Atherosclerosis now has a proven inflammatory mechanism^2^. The mechanism of resolution of atherosclerotic inflammation includes eicosanoids termed specialized pro-resolving mediators (SPMs) such as Resolvin D1 (RvD_1_)^3^. Inflammation resolution is defective in atherosclerosis, and may involve deficiencies of SPMs, phagocytosis of apoptotic leukocytes (efferocytosis) and resolution-phase macrophages (Rm), which have an M2-like phenotype^4,1,5,6,7^.

SPMs may activate adenylate cyclase (AC)-coupled G-Protein Coupled Receptors (GPCRs) to generate cyclic-3’5’-adenosine monophosphate (c-AMP), in a similar manner to PGI_2_, the eicosanoid of vascular homeostasis^3,6,8^. Cyclic-AMP activates transcription factors such as cyclic-AMP response element binding protein 1 (CREB1) and Activating Transcription Factor 1 (ATF1), which modulate gene expression by binding to cyclic-AMP response elements (CRE). Thus, as cyclic-AMP drives both Rm and M2 macrophage phenotypes, and is linked to SPMs and PGI_2_, cyclic-AMP may promote inflammation resolution^9,10^. Moreover, the immediate early cyclic-AMP response genes, *FOS* and *NR4A2*, have been linked to suppression of inflammation^11–13^.

Intraplaque hemorrhage (IPH) is a second key process in advanced atherosclerosis and also promotes intraplaque inflammation and plaque rupture ^14,15^. IPH is associated with a macrophage phenotype, Mhem, that has atheroprotective properties and may reduce the atherogenic potential of IPH^16,17^. Hematoma resolution *in vivo* and Mhem macrophages require macrophage Adenosine Monophosphate Activated Kinase (AMPK) and *ATF1*^18^. In IPH, ATF1 activates a set of genes, notably Heme Oxygenase 1 (*HMOX1,* HO-1), which in concert suppress oxidative stress and cellular lipid overload, isolate iron and lipid from each other, and suppress inflammatory activation^16,17, 19–21^.

Hemin-activated genes are ATF1-dependent, enriched in cyclic-AMP-response elements (CRE), and biochemically ATF1 activates *HMOX1* by binding to a CRE-like half-site at −4200bp. So, ATF1 has been linked to both inflammation resolution and resolution of IPH. CREB1 and ATF1 are very close homologues, and both are cyclic-AMP-responsive and they bind to related sequences. However, we found a curious *in vivo* specificity, as *Atf1* deficiency *in vivo* impaired erythrocyte clearance but not efferocytosis (clearance of apoptotic neutrophils), while si-*Creb1* suppressed efferocytosis^18^.

Current thinking on transcriptional activation distinguishes TFs as lineage-determining (LDTF) or stimulus-dependent (SDTF)^22^. Typically, SDTF bind to lineage-related accessible areas of chromatin, and cause further recruitment of chromatin remodellers, leading to lineage-selective stimulus-dependent activation^22^. Thus in a given stimulus, gene-specific TF recruitment would typically precede chromatin remodeller recruitment and chromatin remodelling^23^.

A key family of chromatin remodellers is Switch / Sucrose non-fermentable (SWI/SNF)) ^22^, and a key mammalian member is SWI/SNF Related Matrix Associated Actin Dependent Regulator of Chromatin, Subfamily A, Member 4 (*SMARCA4*, also known as Brahma related gene 1 (*BRG1*))^24^. It mediates ATP-dependent to nucleosome shuffling to increase accessibility to further cofactors and enzymes, thereby playing a key role in gene activation^24^.

Genome Wide Association (GWAS) studies for atherosclerosis and cardiovascular disease (CVD) have pointed to risk single nucleotide polymorphisms (SNPs) in the vicinity of the *SMARCA4 locus*^25^, with the lead SNP being rs1122608 in intron 29 of *SMARCA4* and rs6511720 in intron 1 of the nearby Low Density Lipoprotein Receptor (*LDLR*) gene. Since *SMARCA4* is only 25kb away from the *LDLR* locus, whether or not it is an independent CVD risk gene is an important question.

Here we present data indicating that *SMARCA4* is an atherosclerotic / CVD risk gene acting independently of *LDLR*. We show data pointing to a novel mechanism, in which early hemin-directed chromatin remodelling and SMARCA4 recruitment modulates the preferred DNA-binding sites of ATF1, favouring gene expression for hematoma-resolution over inflammation-resolution. Modulation by inhibitors of Histone Deacetylase 3 (HDAC3) synergise with both hemin and PGI_2_ to induce target genes. Thus it may be possible to simultaneously enhance both IPH resolution and inflammation resolution in advanced atherosclerosis with HDAC inhibitors.

## Methods

Methods are detailed in the Online Supplement.

**Linkage Disequilibrium** (LD) between pairs of SNP were estimated and LD Plots were generated using the website LDLink https://ldlink.nci.nih.gov/ Gene expression associated with chromosome 19 SNPs was examined using the GTex portal. https://gtexportal.org/home/

***In vitro*** macrophage culture followed our previous methods, for human monocyte derived macrophages (hMDM) and mouse bone marrow macrophages (mBMM)^26^. Written informed consent was obtained from participants, the study protocol conforms to the Declaration of Helsinki and was approved by the Institutions research committee. The hMDM were cultured from normal volunteer blood in 10% autologous serum (AHS) and Iscove’s Modified Dulbecco’s Medium (IMDM). Gene knockdown by si-RNA, and gene expression quantification by RT-qPCR were carried out by previous methods^16,26^. ChIP-qPCR, 3C-qPCR and Co-IP were performed using protocols detailed at length (details in Supplement). In the ChIP, 10^5^ macrophages were stimulated, formaldehyde-fixed, glycine-stopped, nuclei lysed, and chromatin sheared by sonication. Capture was with rabbit monoclonal antibodies or matched rabbit monoclonal IgG control and DNA extraction by phenol-chloroform and ethanol/NaCl precipitation. Analysis was by quantitative polymerase chain reaction (qPCR)^27^.

The 3C-qPCR used a minor modification of published methods, with qPCR primer pairs for enhancer and core promoter (details in Supplement)^28^. RNA-Seq was carried out using RNA from 10^5^ macrophages per well, RNA purification by NEB kit, QC by size and quantity and purity, and sequencing by BGI and analysis Partek (details in Supplement). Biological inferences of differentially expressed genes used a combination of standard multiple approaches (details in Supplement). ELISA was performed using kits from Cayman Chemicals following manufacturer’s instructions. Phagocytosis assays were carried out by a minor modification of existing protocols^29^, in which fluorescent phagocytosis baits were counted in the supernatant as well as the macrophages (details in Supplement). Enzyme assays for Platelet Activating Factor Acetyl Hydrolase (PAF-AH) the product of *PLA2G7*, were carried out by minor modification of established assays, using a colorimetric substrate, based on nitrophenyl-PAF (Merck 810857P) (details in Supplement) ^30–32^. HO-1 activity was carried out by a minor modification of established methods, loading hMDM with hemin then measuring its clearance from the cells (details in Supplement) ^33^.

## Results

### *SMARCA4* and *LDLR* – estimate of linkage disequilibrium

Figure 1A shows the Linkage Disequilibrium (LD) plot across the *SMARCA4–LDLR* locus on chromosome 19. The plot shows strong LD of the GWAS CVD lead SNP rs1122608 located in intron 29 of *SMARCA4*, with 14 other SNPs (R^2^>0.9) and weaker LD with an additional 23 SNPs (R^2^ 0.6-0.8) but weak to no LD with SNPs further upstream and downstream. Peaks of recombination can be seen which break the locus into one block containing the majority of the *LDLR* gene, a second containing the lead *LDLR* SNP (rs6511720), exon 1 of *LDLR* and part of the intergenic region, a third containing the *SMARCA4* lead SNP and the intron and last 4 exons of the gene, and a fourth containing the remainder of the *SMARCA4* gene. The two lead SNPs occur in different LD blocks and are acting independently from each other, with the two SNPs having an R^2^ of 0.2087. We next examined the evidence for association of the two lead SNPs with expression levels of genes using the GTex database. As shown in Supplementary Figure 1, LDLR expression is seen in a wide range of tissues with particularly high levels in adrenals, lung and ovary, and lower levels in liver. *SMARCA4* expression is also seen in a wide range of tissues but with levels being roughly 5-fold lower than for the LDLR. The *SMARCA4* SNP shows significant evidence for expression of *SMARCA4* levels in 10 different tissues, with examples of the pattern of expression shown in Figure 1B and 1C. Interestingly the *LDLR* SNP shows no significant evidence of affecting the expression of LDLR but does show an association with levels of expression of *SMARCA4* (Figure 1D). For both SNPs, expression of *SMARCA4* is lowest in individuals homozygous for the “protective” rare T allele.

**Figure 1.**
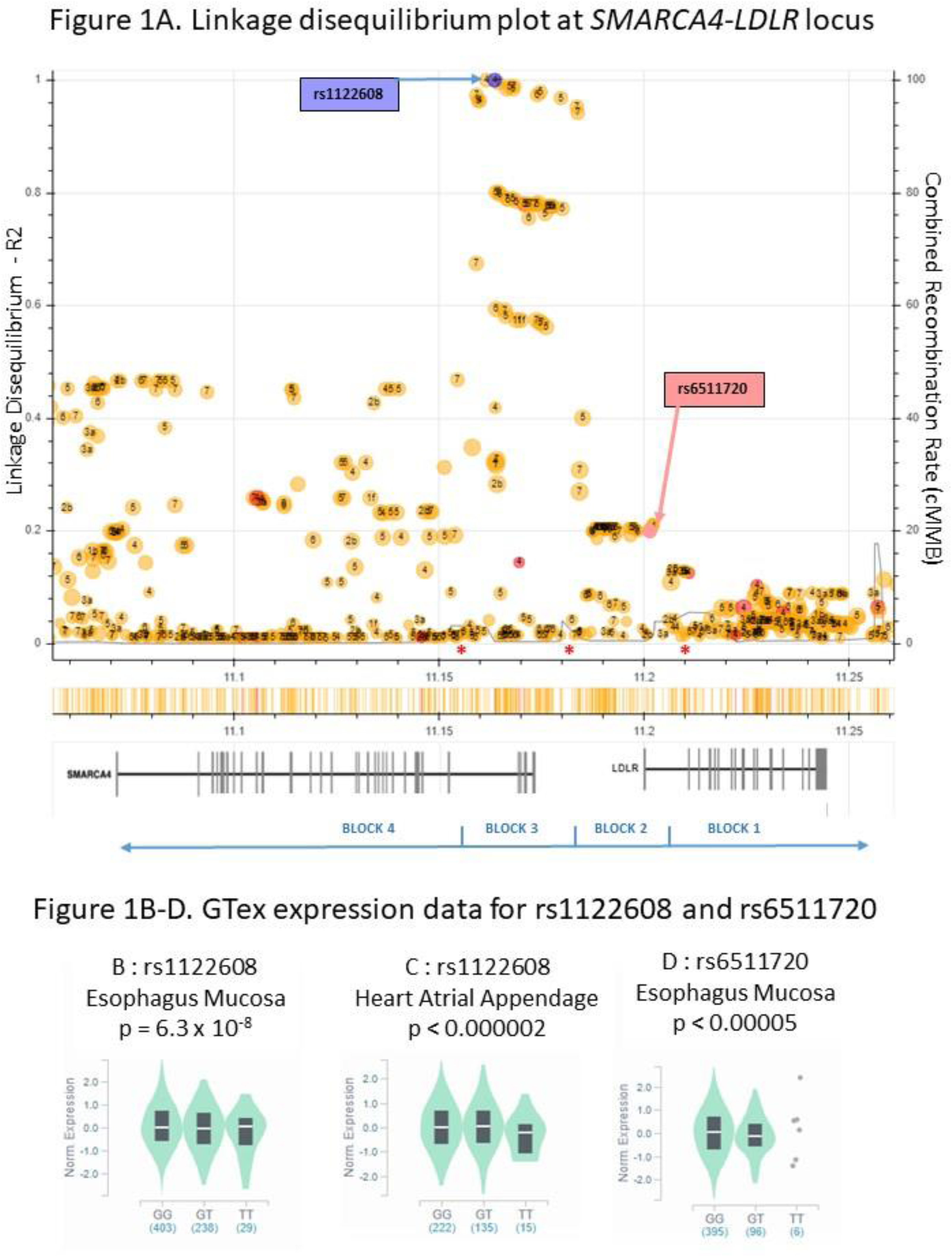
Linkage Disequilibrium (LD) plot and eQTL expression plots for SNPs in the *SMARCA4 – LDLR* locus on chromosome 19. **1A. Linkage Disequilibrium (LD) plot plots for SNPs in the *SMARCA4 – LDLR* locus** The location of the lead *SMARCA4* SNP rs1529729 is shown in blue, and the lead *LDLR* SNP rs6511720 is shown in pink. Each dot indicates the location of the SNP on the chromosome on the X-axis and the extent of LD with rs1529729 on the Y-axis. The numbers in each dot correspond to their regulome DB classification (http://www.regulomedb.org/index). The degree of Linkage Disequilibrium (LD) between rs1529729 and all other SNPs is shown by their height on the left hand Y-axis. An LD of >0.8 would be considered strong LD and between 0.4-0.2 would be considered weak. An LD of below 0.2 indicates essentially no significant LD. Below the plot are shown the location of the *SMARCA4* and *LDLR* genes with vertical bars representing exons. The Y-axis on the right hand side indicates rate of recombination and the line in blues indicates the rate of recombination across the locus. Small peaks of recombination can be seen indicated by *. Data from website LDLink https://ldlink.nci.nih.gov/ **Figure 1B-D. eQTL Violin expression plots for SNPs in the *SMARCA4 – LDLR* locus.** Data from the GTex portal. https://gtexportal.org/home/ Figure 1B. rs1122608 expression with SMARCA4 mRNA in Esophagus Mucosa Figure 1C. rs1122608 expression with SMARCA4 mRNA in Heart Atrial Appendage Mucosa Figure 1D. rs6511720 expression with SMARCA4 mRNA in Esophagus Mucosa

### Specificity of gene expression in response to cyclic-AMP or hemin

Heme induces *HMOX1* via the cyclic-AMP responsive TF p-ATF1 and a cyclic-AMP response element (CRE) half-site at approximately −4.1kb relative to the start codon^16^. To assess its pharmacological potential, whether cyclic-AMP induced *HMOX1* gene expression was tested. Curiously, 100μM di-butyryl-cyclic-AMP (dbc-AMP) (a widely used cell permeable cyclic-AMP analogue at a widely used concentration)^34^ had no effect on *HMOX1* expression by hMDM at any point over a protracted time-course (Figure 2A). Similarly, hemin did not alter expression of classical cyclic-AMP response genes *FOS*, *NR4A2* or *PLA2G7* (Figure 2A and Supplemental Figure 2). In contrast, dbc-AMP induced the cyclic-AMP response *NR4A2* over 20-fold and hemin induced *HMOX1* over 10-fold (Figure 2A). This specificity extended to other genes as dbc-AMP but not hemin also induced *FOS* and *PLA2G7* (Supplemental Figure 2). *PLA2G7* encodes PAF-AH which degrades the inflammatory mediator PAF^35–37^, which was taken further in the functional studies. Effects with dbc-AMP were replicated with 1μM PGI_2_ (Supplemental Figure 3), indicating pathophysiological representativeness.

**Figure 2.**
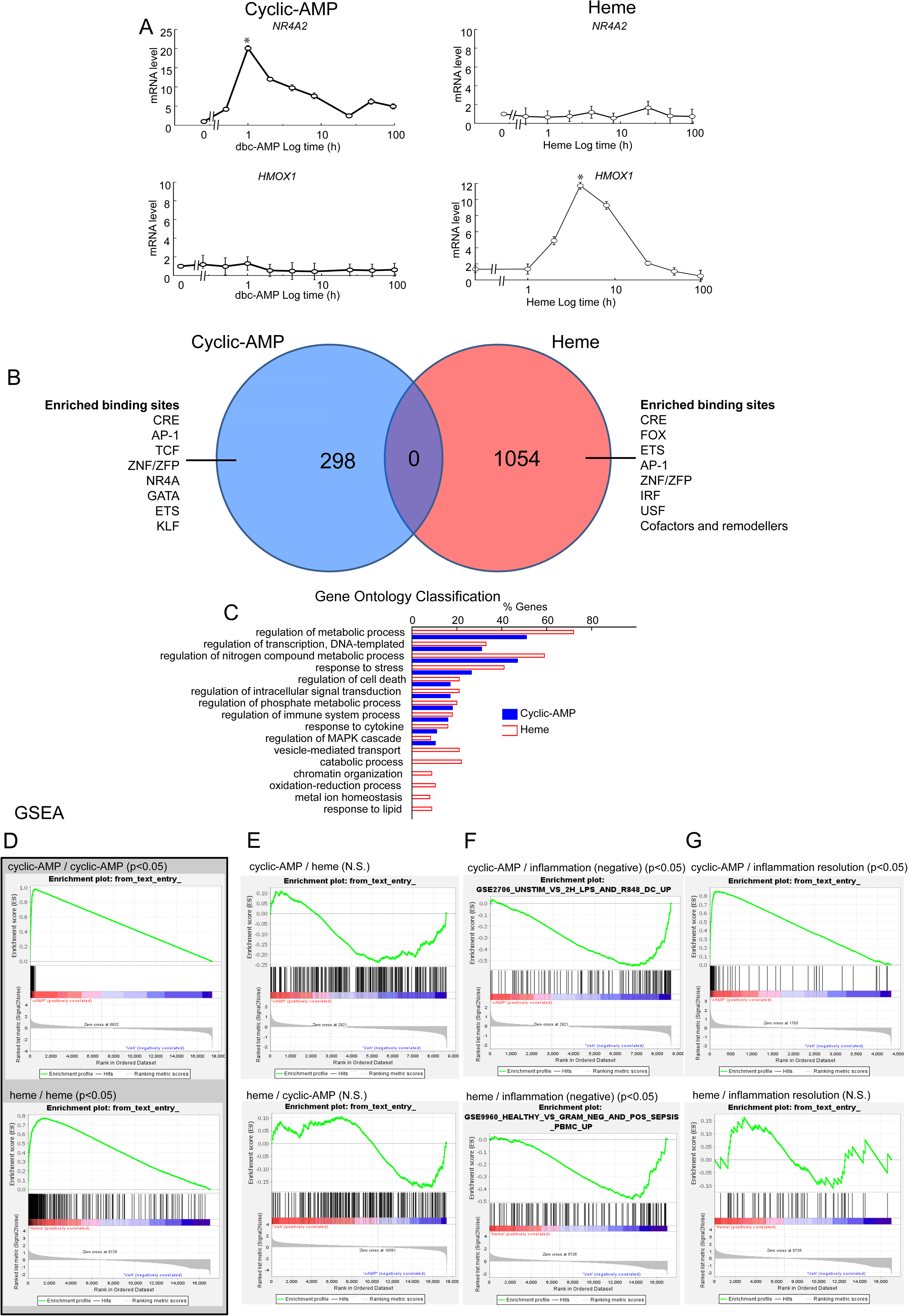
Specificity of gene expression induced by dbc-AMP and hemin. **A, Specificity of responses of *HMOX1* and *NR4A2* to hemin or cyclic-AMP over 168h timecourses by RT-qPCR.** X-axes, time in hours after hemin or cAMP treatment (log scale). Y-axes, induction of indicated genes in fold from baseline, calculated by 2^-ΔΔCt^ method as standard. Data are shown with additional genes in the Supplemental Online. Left– stimulation with 100μM dbc-AMP (n=8 donors). Right – stimulation with 10μM hemin (n=8 donors). Upper– *NR4A2*, Lower – *HMOX1*. **B, Venn diagram** showing lack of overlap between hemin-induced genes (red) and cyclic-AMP-induced genes (blue) (http://www.pangloss.com/seidel/Protocols/venn). Intersection – zero genes in common. To each side are the transcription factor binding sites that were calculated to be overrepresented (Methods). Notably, both gene expression profiles are enriched in cyclic-AMP response elements (CRE). **C, Gene Ontology** comparison of hemin-induced (red bars) and dbc-AMP-induced (blue bars) gene expression profiles (http://www.pantherdb.org/). X-axis, % of genes in each GO category (may add up to over 100% if genes fall into more than 1 category). Y-axis, GO categories (selected for being informative). With the exception of genes for catabolism, chromatin organization, redox, metal ion homeostasis and lipids, the GO categories were similar. **D-G, Geneset Enrichment Analysis (GSEA)** to compare hemin-regulated and dbc-AMP-regulated gene expression profiles (www.gsea-msigdb.org). Upper series – cyclic-AMP (as indicated). Lower series - hemin (as indicated). A rapid rise of the green curve to a high peak, clustering of gene hits (barcode) to one end, indicate a good match. Significances are as given. C, leftmost grey background – comparison of gene expression profiles to themselves showing what would be expected with a very close match. D, second panel, there was no significant similarity between hemin and cyclic-AMP. E, third panels, both hemin and cyclic-AMP were inversely related to human inflammation genesets *in vivo*. F, rightmost panels, cyclic-AMP matched an inflammation resolution geneset but hemin did not.

Transcriptomics of hemin and dbc-AMP treated hMDM were next compared to assess this specificity across the genome (Methods). These were on identically prepared hMDM at 24h from isolation, adherence-purified and in 10% AHS / IMDM. Consistency of gene expression data was ensured using standardized HGNC-approved gene names.

When analysed by Venn diagram, (Figure 2B) dbc-AMP regulated 298 genes, hemin regulated 1054 genes, but there was zero overlap between hemin and dbc-AMP modulated gene sets.

Next, a transcription factor binding site (TFBS) analysis was carried out to compare their likely drivers (Methods). The top 8 scoring TFBS for cyclic-AMP (ordered by rank, Figure 2B) were cyclic-AMP Response elements (CRE), AP-1, TCF, ZNF, NR4A, GATA, ETS, KLF. Notably, CRE were the top TFBS for both genesets. Many TFBS in the top 8 were shared between gene sets, indicating that unique combinations of TFBS were unlikely to explain the specificity. There was a greater enrichment in predicted binding sites for chromatin remodelers in the hemin-regulated genes (Figure 2B), such as *SMARCA5*, *NCOA3*, *EP300*, *PRDM4*. Moreover, the hemin-regulated genes included chromatin remodelers such as *SMARCD1* (704 genes) more than the cyclic-AMP-regulated genes (8 genes). The hemin-induced and cyclic-AMP induced gene expression profiles were next compared using Gene Ontology (GO) (Figure 2C). This revealed a broad overall functional similarity between the cyclic-AMP and hemin-regulated gene expression profiles. Hemin-regulated genes matched more highly for chromatin organisation, catabolism, redox, lipid metabolism, metal ion homeostasis.

Geneset Enrichment Analysis (GSEA) allows a more precise quantification than binary name matching and was used as described^38^ (Figure 2D-G). As controls, hemin and dbc-AMP had high self-similarity cores (Figure 2D). When hemin and dbc-AMP were compared with each other, enrichment scores were low and non-significant (Figure 2E), consistent with the findings in Figure 2B. Comparing hemin and dbc-AMP with public datasets showed significant highly negative enrichment scores for inflammation involving human PBMCs (Figure 2F), a pattern found across multiple GSEA datasets on human inflammation (https://www.gsea-msigdb.org/gsea/msigdb/search.jsp). Thus both pathways were anti-inflammatory. There is not yet an established ‘inflammation resolution’ public dataset based on experimental human data, and so a nearest approximation was created from existing data sources including literature and murine-omics (Methods). This approach showed that the dbc-AMP-induced gene expression profile matched to inflammation resolution and that hemin-induced expression profile did not (Figure 2G).

The network architecture of the response were estimated and compared using the STRING estimated STRING database of protein-protein interactions (https://string-db.org/)^39–41^ (Supplemental Figure 4), showing potential large differences between the network topologies, that appeared to involve chromatin remodellers.

Taken together, these results indicated that hemin / dbc-AMP specificity extended beyond key exemplar genes to alternative gene expression programmes with distinctive functional consequences. We postulated that hemin-activated CRE-mediated genes reflected adaptation to erythrocyte clearance and cyclic-AMP-activated CRE-mediated genes reflected adaptation to leukocyte clearance.

### SMARCA4 switches pATF1 gene binding and consequent expression

To explore whether the specificity related to differences in chromatin remodeling, enhancer-promoter gene looping was measured by chromatin conformation capture analysis (3C) and quantitative PCR for the *HMOX1* enhancer^28,42^ (Supplemental Figure 5). Gene looping (Supplemental Figure 5B) and SMARCA4 recruitment (Supplemental Figure 5C) peaked at 1h, which was well before binding of p-ATF1 (Supplemental Figure 5D). This suggested early stimulus-directed recruitment of SMARCA4 to *HMOX1* prior to arrival of p-ATF1, which was curious, as usually TF binding leads to chromatin remodelling, rather than the other way around.

Because SMARCA4 bound to *HMOX1* prior to p-ATF1, it was postulated that SMARCA4-deficiency would alter recruitment of p-ATF1 to *HMOX1* and this was tested in hMDM using si-RNA. Knockdown of SMARCA4 with si-*SMARCA4* suppressed p-ATF1 binding to the *HMOX1*-enhancer −4100CRE site (Figure 3A), while it increased p-ATF1 binding to CRE-sites of the cyclic-AMP responders *NR4A2* and *FOS* (Figure 3B-C). Si-SMARCA4 abrogated 3C signal by qPCR, indicating that the early hemin-induced *HMOX1* enhancer-promoter loops were SMARCA4-dependent (Figure 3D).

**Figure 3.**
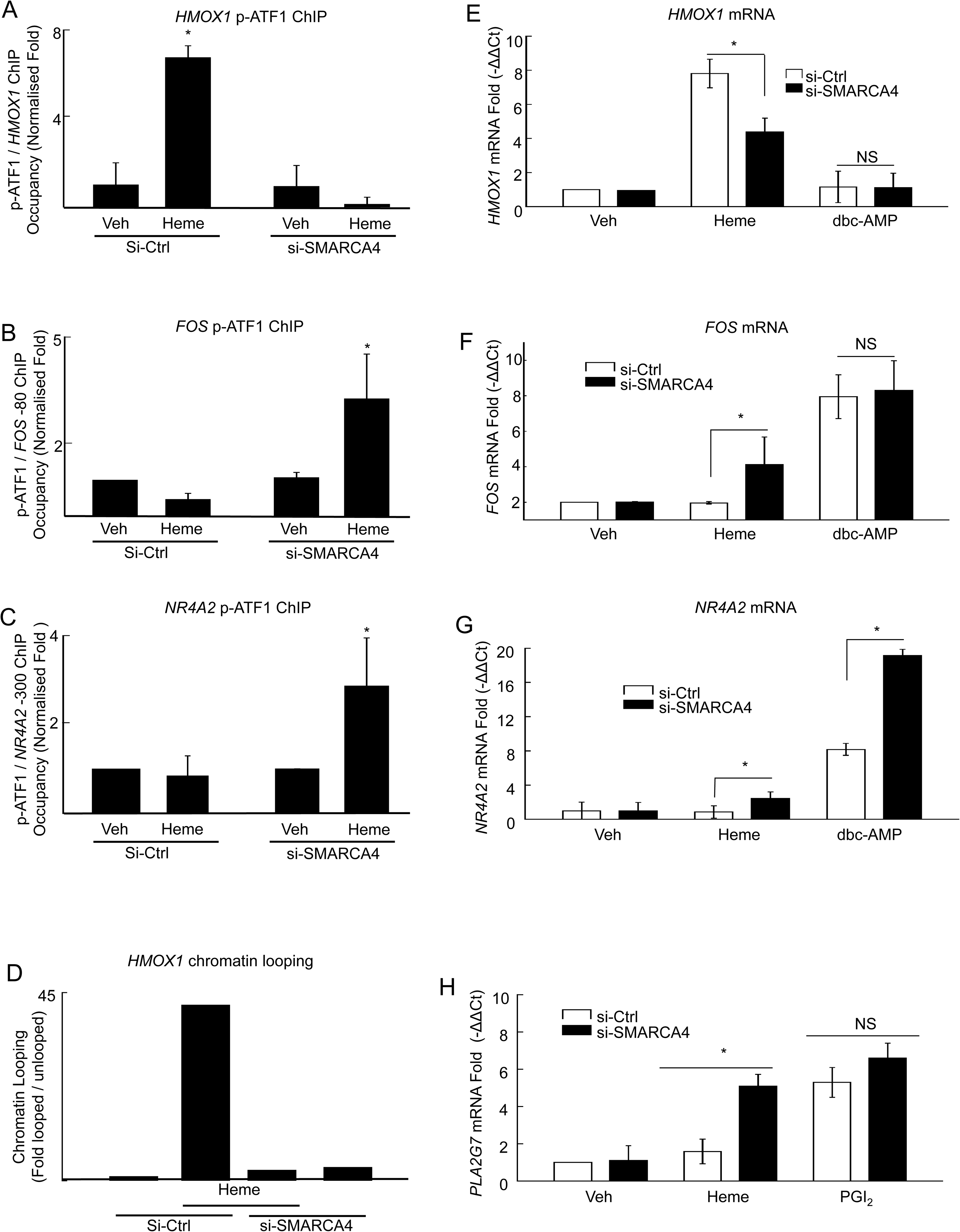
SMARCA4-knockdown shifts p-ATF1 from *HMOX1* to *FOS* and *NR4A2*, and functionally redirects hemin-responses to resemble PGI_2_ responses. **A-C, effects of gene knockdown of SMARCA4 on localisation of p-ATF1** on genes for *HMOX1*, *NR4A2* or *FOS*. In each, knockdown was by transfecting human blood-derived macrophages with si-RNA for SMARCA4 (Dharmacon) using Interferin (Polyethyleneimine, Polysciences). ChIP and RT-qPCR were as before (detailed Methods). Data are mean ± SE, n=7 donors, *p<0.05, ANOVA. **D, effect of SMARCA4 knockdown on chromatin looping.** Y-axis, chromatin looping by 3C, fold relative to baseline. X-axis, treatment. N=2. **E-G, effects of gene knockdown of SMARCA4 on gene expression for *HMOX1*, *FOS* and *NR4A2* in response to hemin or dbc-AMP (by RT-qPCR).** E-G all show RT-qPCR on the same set of cells with or without si-*SMARCA4* (or non-targeting control) and with or without or dbc-AMP or hemin. Y-axes, *HMOX1* (E); *FOS* (F), *NR4A2* (G). X-axes, treatments as indicated. As in other experiments dbc-AMP was 100µM and hemin 10µM. Control non-targeting siRNA (open bars) SMARCA4-siRNA (closed) as indicated. Data are mean ± SE, n=7 donors, *p<0.05, Wilcoxon. **H, in human blood-derived macrophages, SMARCA4-knockdown causes hemin to induce PLA2G7 in a similar manner to PGI_2_.** Y-axis, *PLA2G7* gene induction, fold from baseline, measured by −2^ΔΔCt^. X-axis, treatments. PGI_2_, 100nmol.L^-1^ (100nM) Iloprost for 4h. Control si-RNA (open) and si-*SMARCA4* (filled) as indicated and as other panels. Data mean ± SE, *p<0.05, n=5, Wilcoxon.

That left a question of whether the redirected p-ATF1 binding had a functional role in gene expression. To answer this, RT-qPCR showed that si-*SMARCA4* suppressed hemin-induced *HMOX1* (Figure 3E) and caused hemin to instead induce *FOS* and *NR4A2* (Figure 3F, G). *SMARCA4*-knockdown did not affect cyclic-AMP-induced *FOS* and *NR4A2,* excluding general or non-specific effects on transcription (Figure 3F, G). Therefore, the effect of SMARCA4 on p-ATF1-binding results in alternative gene expression, with SMARCA4 favouring hemin induction of *HMOX1* over *FOS* and *NR4A2*.

There is a paucity of data on gene effectors of inflammation-resolution. Expression of *PLA2G7* (PAF-AH) was studied in response to PGI_2_ and hemin for a better understanding of pathophysiological significance. In hMDM, hemin-stimulation did not induce *PLA2G7* under control conditions, but on si-*SMARCA4-*mediated knockdown, hemin treatment strongly induced *PLA2G7* (Figure 3H). This indicated that SMARCA4-deficiency caused hemin-induced gene responses to resemble prostacyclin responses. As PAF is atherogenic and PAF-AH is atheroprotective, this is likely to have functional significance.

### HDAC3 inhibition and pathophysiological responses

The next question was linkage to pathophysiology and pharmacology. The connection between cyclic-AMP and inflammation resolution is not well understood, but *FOS* and *NR4A2* may limit inflammation and are cyclic-AMP-inducible. *FOS* and *NR4A2* were measured in response to PGI_2_, or specialised mediators of inflammation resolution e.g. RvD_1_. Apremilast, is a PDE4 inhibitor, selectively increases cyclic-AMP and is clinically licensed as an anti-inflammatory drug^43^. PGI_2_ and RvD_1_ induced *FOS* and *NR4A2* (Supplemental Figure 6), with greater expression in the presence of Apremilast, consistent with mediation by cyclic-AMP (Supplemental Figure 6).

Histone deacetylase inhibitors are entering clinical use and could potentially facilitate activation of pro-resolving genes. This was addressed for hemin-induced *HMOX1*. The histone acetyl modification (Histone H3 Lys9 acetate H3K9Ac) was found on *HMOX1* at 1h after hemin stimulation (Figure 4A). The broad-spectrum HDAC inhibitor Trichostatin-A increased recruitment of SMARCA4 to *HMOX1* (Figure 4B) and increased *HMOX1* mRNA (Figure 4C). This indicates that hemin drives *HMOX1* histone acetylation early after hemin stimulation, which recruits SMARCA4 (as increasing histone acetylation increases SMARCA4 recruitment) and this results in increased *HMOX1*. Next, we assessed the HDACi RGFP966, at a concentration at which it is a selective inhibitor for HDAC3 (100nM) (Figure 4D-E). PGI_2_ and hemin were maintained as standard stimuli, and *HMOX1* and *NR4A2* as more specific target genes (than *FOS*). Alone, RGFP966 had no significant effect on either *HMOX1* or *NR4A2*. As before, hemin-induced *HMOX1* and PGI_2_ induced *NR4A2*. Although it had no effect on its own, RGFP966 increased both hemin-induced *HMOX1* and PGI_2_-induced *NR4A2* (Figure 4D-E). This raises the possibility that a selective HDAC3-inhibitor might promote both inflammation-resolution and IPH-resolution in advanced atherosclerotic plaques.

**Figure 4.**
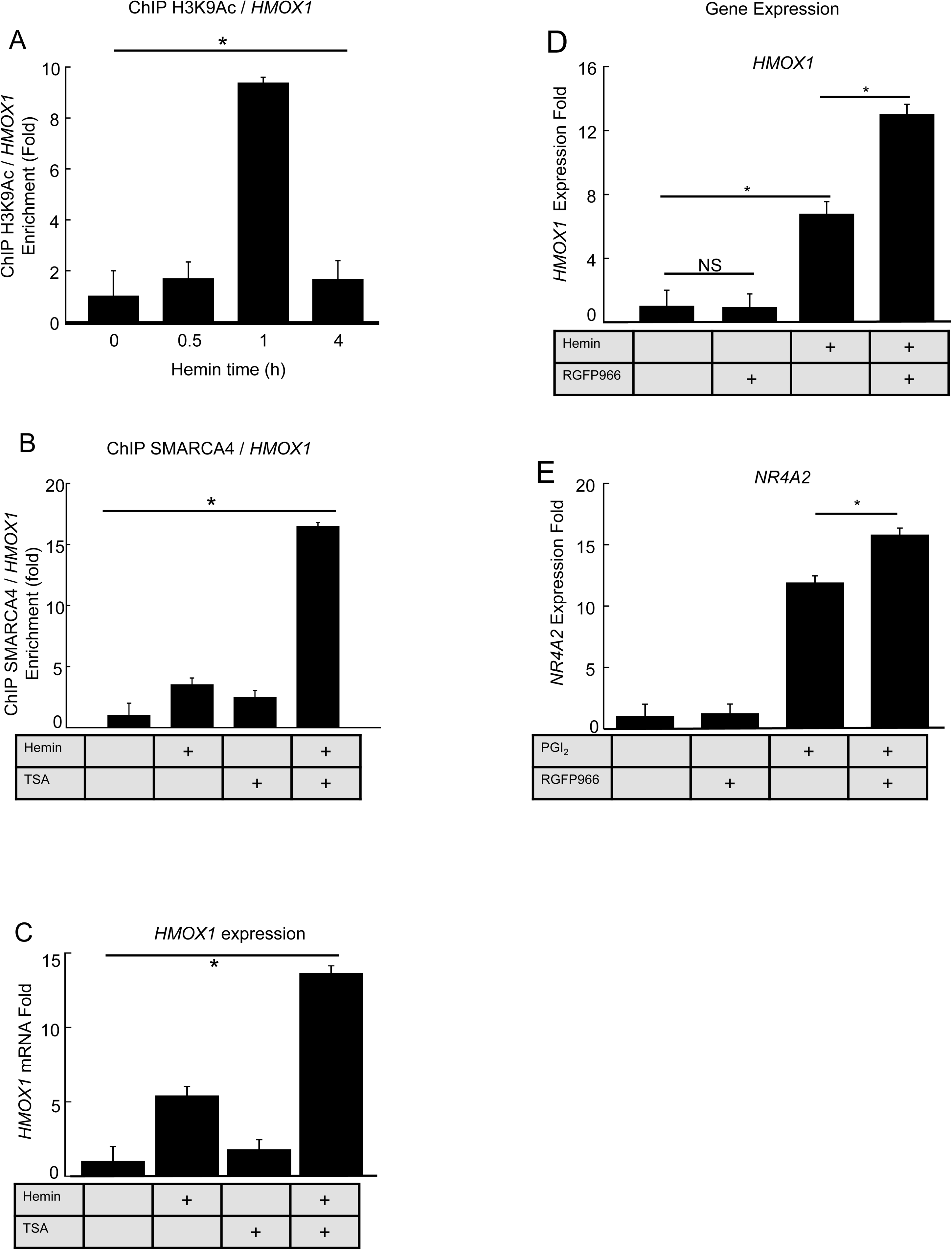
SMARCA4 is itself recruited by histone acetylation. **A, Histone acetylation of Histone H3 Lysine 9 (H3K9ac) at the *HMOX1*-4100 site**, by ChIP-qPCR in human blood-derived macrophages, as for the other Figures. Data are mean ± SE, n=5 donors, *p<0.05, ANOVA. **B, Recruitment of SMARCA4 to the *HMOX1-4100* site with and without 100nM Trichostatin-A** by ChIP-qPCR in human blood-derived macrophages, methods as before. Trichostatin-A (100nmol.L^-1^, 100nM) was added 20 minutes before hemin and macrophages incubated for 1h. Data are mean ± SE, n=5 donors, *p<0.05, ANOVA on ranks. **C, Hemin-induced *HMOX1* expression by hMDM, with and without 100nM Trichostatin-A**, by RT-qPCR in human blood-derived macrophages, methods as before. Y-axis, *HMOX1* Fold expression by RT-qPCR and −2^ΔΔCt^ method, X-axis, culture conditions reflecting 4h stimulation with 10μM hemin with or without 100nM Trichostatin-A. Data are mean ± SE, n=5 donors, *p<0.01, Repeated measures ANOVA (also *p<0.001 Kruskal Wallis). **D, Hemin-induced *HMOX1* expression by hMDM, with and without 100nM RGFP966** by RT-qPCR in human blood-derived macrophages, methods as before.Y-axis, *HMOX1* Fold expression by RT-qPCR and −2^ΔΔCt^ method, X-axis, culture conditions reflecting 4h stimulation with 10μM hemin with or without 100nM RFGP966. Data are mean ± SE, n=5 donors.*p<0.01, Repeated measures ANOVA (also *p<0.001 Kruskal Wallis). **E, PGI_2_-induced *NR4A2* expression by hMDM, with and without 100nM RGFP966** by RT-qPCR in human blood-derived macrophages, methods as before. Y-axis, *NR4A2* Fold expression by RT-qPCR and −2^ΔΔCt^ method, X-axis, culture conditions reflecting 4h stimulation with 1μM PGI_2_ (as Iloprost) with or without 100nM RFGP966. Data are mean ± SE, n=5 donors. *p<0.01, Repeated measures ANOVA (also *p<0.001 Kruskal Wallis).

### Pathophysiology: SMARCA4 switches gene activation from genes for erythrocyte disposal to genes for disposal of inflammatory mediators and apoptotic leukocytes

It was next postulated that SMARCA4 redirected ATF1 from genes oriented to leukocyte disposal to those oriented to erythrocyte disposal. That way, loss of SMARCA4 would ‘rewire’ macrophages so that hemin-responses would instead drive leukocyte-resolution effectors.

Macrophages internalise debris to be cleared, such as apoptotic leukocytes at the end of inflammation, or oxidatively damaged erythrocytes. This process has at least 2 components, internalisation and then degradation of the internalized material. The internalisation step is intensively studied, often measured as the phagocytic index (PI). The degradation step is less studied but just as important and relates to catabolic enzymes (Figure 5A). Importantly, the PI measurement (phagocytosed objects counted divided by macrophages) is *decreased* by enzymatic digestion of the phagocytosed material. Thus, when phagocytosis and accompanying catabolism are coregulated, these are potentially confounded and require unmixing.

**Figure 5.**
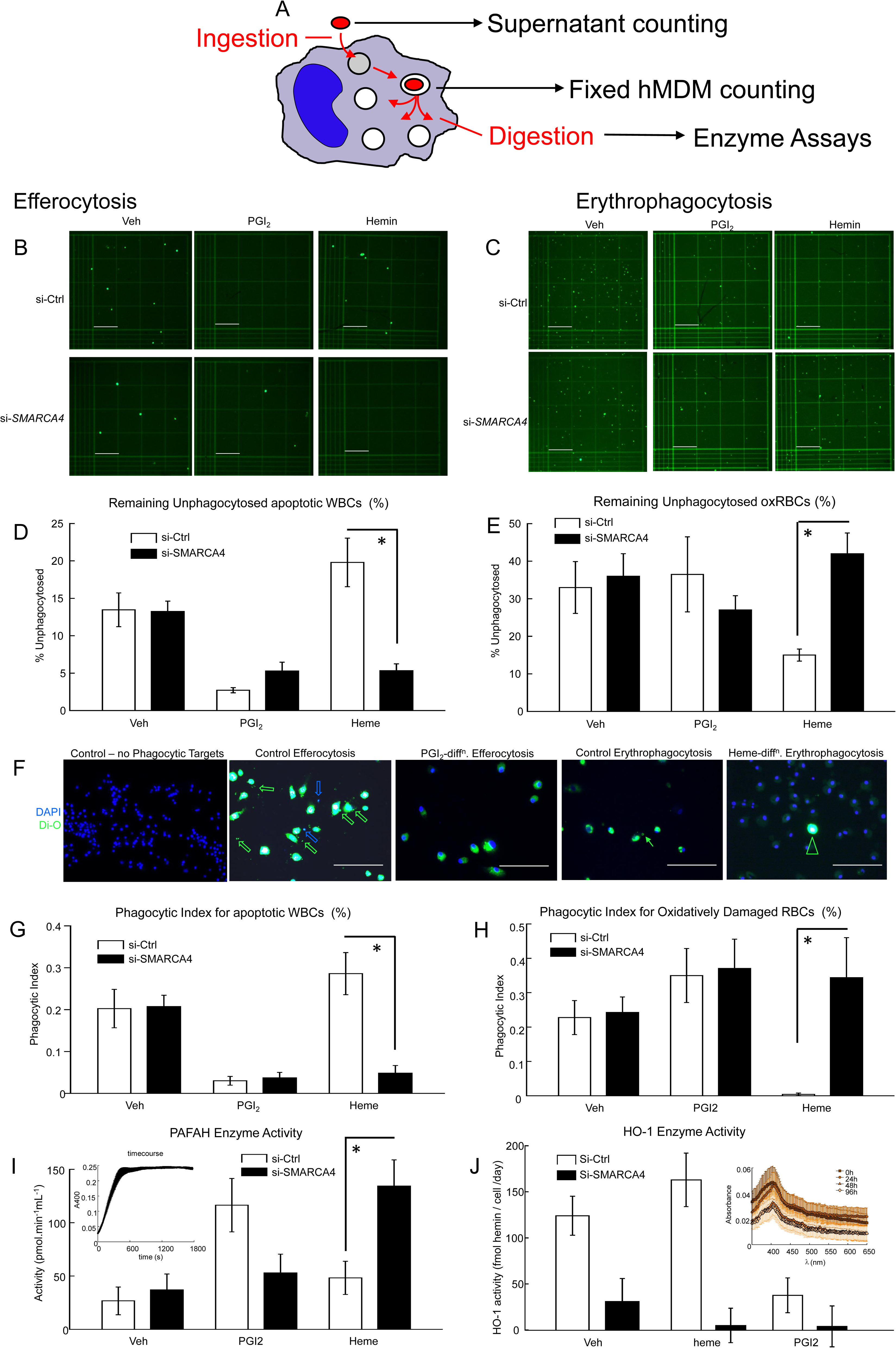
Functional significance of SMARCA4-directed gene expression. **A, Schema of extended phagocytosis scheme (internalisation and then degradation).** Estimation of cytoplasmic phagocytosed cells, as commonly evaluated, is a resultant of internalisation and metabolic clearance of the internalized material. The unphagocytosed bait in the supernatant was evaluated as well as the internalized bait. Degradation post-phagocytosis is a reflection of enzyme activity. **B, Efferocytosis, residual supernatant unphagocytosed apoptotic WBCs.** RAW264.7 white blood cells were killed by UV-induced apoptosis and then added to cultures of hMDM. Images are representative of n=5 donors (quantified in D). Images were taken by adding cell culture supernatant to a hemacytometer photographing with a fluorescence microscope. Treatment conditions as indicated (Methods Online). *p<0.05, n=5 Wilcoxon, si-SMARCA4 suppresses unphagocytosed supernatant apoptotic WBCs, indicating that it promotes their uptake **C, Erythrophagocytosis, supernatant unphagocytosed RBCs.** Autologous erythrocytes were oxidatively damaged by hydrogen peroxide and then added to cultures of hMDM. Images are representative of n=5 donors (quantified in E). Images were taken by adding cell culture supernatant to a hemacytometer photographing with a fluorescence microscope. Treatment conditions as indicated (Methods Online). *p<0.05, si-*SMARCA4* increases unphagocytosed supernatant oxidized RBCs relative to non-targetting control, indicating that it promotes their uptake. **D, Quantification of efferocytosis, residual supernatant unphagocytosed WBCs.** Y-axis, % of dead white blood cells left unphagocytosed. X-axis, treatment. **E, Quantification of erythrophagocytosis, residual supernatant unphagocytosed RBCs.** Y-axis, % of damaged red blood cells left unphagocytosed. X-axis, treatment. **F, Images of internalized bait within macrophages at 18h.** Representative images, n=5 donors. Scalebar – 100μm. Blue, nuclei (DAPI), green – DiO label. Open blue arrows, nuclei. Open green arrows, macrophages containing phagocytosed di-O-labelled dead WBCs. Fine green arrow, Di-O-labelled erythrocyte within a macrophage. Open green arrowhead, macrophage containing abundant DiO derived from RBCs but no identifiable RBCs. Notably, internalized WBCs are less conspicuous after PGI_2_ treatment and internalized RBCs are less conspicuous after hemin pretreatment. Quantified in G-H **G, Phagocytic index for apoptotic WBCs.** Y-axis, phagocytic index as conventionally calculated (phagocytosed apoptotic WBCs / macrophages). X-axis, treatment. *p<0.05, n=5 Wilcoxon, si-*SMARCA4* suppresses the phagocytic index. **H, Phagocytic index for ox-RBCs.** Y-axis, phagocytic index as conventionally calculated (phagocytosed ox-RBCs / macrophages). X-axis, treatment. *p<0.05, n=5 Wilcoxon, si-*SMARCA4* suppresses the phagocytic index relative to non-targetting siRNA. **I, PAFAH enzyme activity.** Inset, enzyme kinetic; x-axis, time; y-axis, absorbance from coloured product. Main, y-axis, enzyme activity (arbitrary units); x-axis, treatment. *p<0.05, n=5 Wilcoxon, si-*SMARCA4* enables hemin to increase PAFAH activity relative to non-targetting control. **J, HO-1 enzyme activity.** Inset, enzyme kinetic; x-axis, wavelength (λ)(nm); y-axis, absorbance from coloured product. An absorbance peak specific to hemin is seen (Soret) and decreases with time. Main figure, y-axis, enzyme activity (fmol hemin / cell per day); x-axis, treatment. *p<0.05, n=5 Wilcoxon, si-*SMARCA4* abolishes ability of hemin to increase HO-1 activity.

Apoptotic WBCs and oxidatively-damaged RBCs are both classic models of phagocytosis^44–46^. Apoptotic WBCs and OxRBC that were left in the culture supernatant after 18h of incubation were counted in addition to phagocytic index (Figure 5B-E). This was to evaluate an overall measure of clearance, i.e. debris lost to the extracellular space.

Unphagocytosed supernatant fluorescent WBC and RBC were counted by hemacytometer (representative photomicrographs in Figure 5B-C). Prostacyclin decreased uncleared WBCs (Figure 5D), hemin decreased uncleared RBCs (Figure 5E), but not *vice versa* (Figure 5D-E). SMARCA4-si-RNA caused hemin to decrease supernatant apoptotic WBCs (Figure 5E), similar to prostacyclin. Conversely, SMARCA4-si-RNA abolished hemin-mediated suppression of uncleared RBCs (Figure 5E), indicating that SMARCA4 promoted erythrocyte phagocytosis.

Next, the classical phagocytic index for WBCs and RBCs was calculated on adherent macrophages in the same wells (after fixation and counterstaining) (Figure 5F-H). Representative micrographs are shown in Figure 5F, and quantification is in Figure 5G-H. Prostacyclin decreased observable phagocytosed apoptotic WBCs (Figure 5G). Hemin decreased observable phagocytosed ox-RBCs (Figure 5H). Decreased phagocytosis was excluded as a cause, as extracellular apoptotic WBCs and ox-RBCs had also been correspondingly decreased (Figure 5D-E). Therefore, the effects related to increased intracellular degradation, respectively of apoptotic WBCs by prostacyclin and of ox-RBCs by hemin. SMARCA4-si-RNA abolished the ability of hemin to suppress intramacrophage observed ox-RBCs (Figure 5H), and enabled the ability of hemin to suppress intramacrophage observed apoptotic WBCs (Figure 5G).

Next, specific enzyme activities were measured. The enzyme activities of HO-1 and PAF-AH were measured according to published methods (Figure 5I-J) ^30–33^. Under control conditions, PGI_2_ elevated PAF-AH activity, whilst hemin did not (Figure 5I). Conversely, hemin increased HO-1 activity but PGI_2_ did not (Figure 5J). SMARCA4-si-RNA altered the hemin response to increase PAF-AH activity and decrease HO-1 activity (Figure 5I-J).

Thus, taken together, this set of experiments indicated that hemin promotes erythrocyte clearance at the expense of inflammation resolution, and this is countered by gene knockdown of SMARCA4.

### Results summary

Taken together, these data indicate: (1) *SMARCA4* and *LDLR* are independent risk loci for cardiovascular disease; (2) cyclic-AMP and hemin induce distinct gene expression responses via CRE-sites; (3) hemin initiates chromatin remodelling with SMARCA4 recruitment prior to ATF1 recruitment; (4) SMARCA4 diverts ATF1 from cyclic-AMP responsive genes to hemin-responding genes; (5) SMARCA4 knockdown functionally converts the hemin response to mimic PGI_2_, including preferential disposal of leukocytes and inflammatory mediators over erythrocytes and hemin; (6) HDAC inhibition modulates SMARCA4 recruitment and HDAC3 inhibition synergises with both hemin and PGI2. Thus HDAC3 inhibitors may be beneficial in advanced atherosclerotic plaques characterised by both IPH and impaired inflammation-resolution.

## Discussion

This work defines a novel role for *SMARCA4* in human blood-derived macrophages, switching them from leukocyte disposal to erythrocyte disposal, two important processes in atherosclerosis, and shows that it is a clinical cardiovascular risk locus genetically independent of *LDLR*. It is normally considered that TF-gene binding drives chromatin remodelling. However, we show here for the first time that chromatin remodelling may drive the specificity of TF-gene binding. Thus, SMARCA4 promotes activation of the hemin-response gene, *HMOX1,* over cyclic-AMP-response genes which functionally mediates a switch from leukocyte disposal to erythrocyte disposal. This points to reciprocity between inflammation-resolution and hematoma-resolution, both important processes in atherosclerosis.

The cyclic-AMP/CREB pathway is well-characterized and exemplifies SDTF-mediated gene activation^22,23, 47–50^. Reviews typically frame this sequence as primary activation of a TF, causing binding to a specific DNA sequence determined only by the sequence, then recruitment of chromatin remodellers, then of RNA Polymerase. CREB1 is activated by phosphorylation by PKA downstream of cyclic-AMP^51^. Phospho-(activated) CREB1, binds cyclic-AMP response elements in gene regulatory sequences, then recruits CBP/p300, which acetylates Histone H3 at several lysine residues^47^. Acetylated histones recruit chromatin remodelers, such as SWI/SNF complexes (including SMARCA4) that contribute to architectural changes such as chromatin opening and chromatin looping, leading to assembly of RNA polymerase at the transcriptional start site, which initiates transcription^23^.

The data presented here indicate a novel mechanism in which hemin-driven chromatin remodelling that pivots on SMARCA4 favours the *HMOX1* CRE for p-ATF1, preferentially inducing *HMOX1*. There were two histone modifications identified, although there may be many more. The overall functional effect is to reroute hemin-stimulated gene activation from inflammation-related effectors (e.g. *PLA2G7*) to erythrocyte-related effectors (e.g. *HMOX1*) and RBC disposal to WBC disposal.

Figure 6 shows a schema for this interaction. We propose that signals from hemin cause SMARCA4 recruitment to *HMOX1*, causing gene looping and preferential recruitment of p-ATF1 once it is activated. It is uncertain why this site is preferential, but it may reflect greater chromatin open-ness or direct protein-protein binding. The SMARCA4 recruitment is likely due to histone acetylation at the hemin-sites and histone phosphorylation at the cyclic-AMP sites. The overall effect enables stimulus-specific chromatin remodeling that switches the transcriptional response from cyclic-AMP to hemin even with almost identical TFs and TFBS. This retunes pathophysiological responses from inflammation-resolution to hematoma-resolution. *PLA2G7* was identified as a downstream effector of cyclic-AMP-mediated and prostacyclin-mediated stimulation. SMARCA4 suppression enabled hemin-mediated *PLA2G7* induction. The enzyme encoded by *PLA2G7* is known both as an isoform of PAF-AH and an isoform of secretory phospholipase 2 (s-PLA_2_). Therefore, *PLA2G7* is known for two opposing roles. As s-PLA_2_, PLA2G7 may degrade LDL to an atherogenic form^52^. As PAF-AH, it degrades the atherogenic inflammatory mediator PAF^37,53–55^. PLA2G7 is a target of the drug darapladib, which failed in Phase III clinical trials of vascular disease prevention^52^. Consistent with this, *PLA2G7* has no net effect in Mendelian randomization studies^56^. Therapeutic strategies that differentiate between these 2 roles may be helpful.

**Figure 6.**
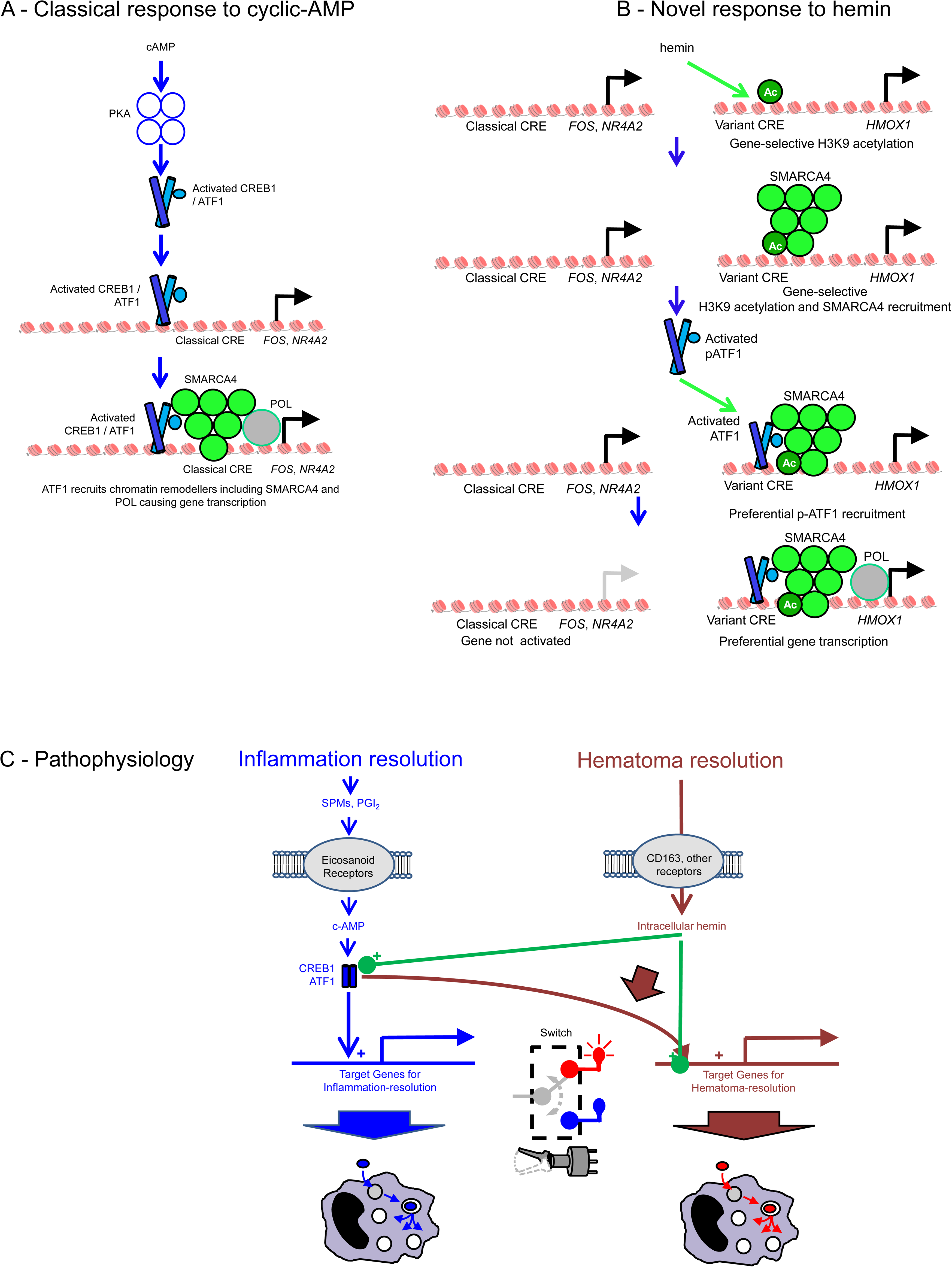
Simplified schema of specificity mechanism. **A, Classical mechanism of gene activation via CRE by cyclic-AMP.** Cyclic-AMP stimulates PKA, which activates ATF1 and CREB1 (crossed cylinders) by phosphorylation. These bind to classical CRE-sites on classically cyclic-AMP-responsive genes (*FOS*, *NR4A2*). The activated TFs on the site then recruit chromatin remodelers (green), and eventually RNA Polymerase (POL, RNAP, grey ovoid, green edge) which transcribes downstream genes eg *FOS*, *NR4A2*. **B, novel variant mechanism, adapted to respond to hemin and with chromatin events occurring first and fine-tuning response specificity.** The first events are hemin-induced histone acetylation of chromatin, with binding of SMARCA4 recruitment. This is before ATF1 is activated. The SMARCA4-bound chromatin then preferentially attracts ATF1 once activated (crossed cylinders), which then recruits RNA Polymerase (POL, RNAP, grey ovoid, green edge) which transcribes an alternative set of downstream genes eg *HMOX*1. **C, Overall pathophysiological significance.** This work points to a relationship between 2 distinct ‘M2-like’ phenotypes that are pro-resolving and anti-inflammatory. Both connect extracellular signals to gene activation via related TFs and TFBS. The cyclic-AMP mediated pathway is likely to come into play in responses to eicosanoid receptors involved in inflammation-resolution. Targets such as NR4A2 and FOS may play roles in inducing clearance molecules and suppressing inflammation. In the novel hemin pathway, hemin-mediated chromatin changes switch over ATF1-mediated gene activation much like a 2-way electrical toggle switch may switch current from one bulb (circuit) or another (centre). This switching is dependent on SWITCH gene SMARCA4, which makes hemin-response genes a preferred target. This may reflect either affinity, or physical opening and closing of the regulatory regions of the genes. The net effect is to generate two alternative related phenotypes of macrophage that are specialised for cleaning up after inflammation; or for cleaning up after a tissue haemorrhage.

Use of ChIP-Seq, rather than ChIP-qPCR as an outcome would extend these data across the whole genome and allow us to make a statement about this mechanism genuinely modulating the p-ATF1 cistrome (set of genome-wide binding sites). At the other end of integration, further application of knockout approaches si-RNA-mediated knockdown and drug inhibitors to hematoma-clearance and inflammation-clearance models in vivo would lend considerable causal pathophysiological relevance. Demonstration of these pathways in human atherosclerotic plaques would also indicate pathophysiological importance. Putting all 3 approaches together in an *in vivo* ChIP-Seq would be extremely powerful.

### Atherosclerosis pathophysiology and switching by SMARCA4

It has become recently understood that impaired efferocytosis and deficiency in SPMs are important features maintaining chronic inflammation in atherosclerosis^1,3^. It would, in principle, be therapeutic to upregulate these processes. It has also become appreciated that IPH promotes inflammation and oxidative stress, promoting atherogenesis^57^. Mhem express the protective enzyme HO-1, the anti-inflammatory cytokine IL-10, have an anti-inflammatory gene profile and are resistant to lipid overload and have less oxidative stress^16,17^. The key gene, *HMOX1*, is induced via AMPK and ATF1, as well as NFE2L2 (Nrf2) in humans and mouse, and deficiency prevents subcutaneous hematoma resolution. Thus, it appears that *Atf1 / Hmox1* - driven Mhem protect against IPH. We show here that macrophage responses to IPH and inflammation resolution have some similarities, but due to SMARCA4 show reciprocity as the SMARCA4 makes hemin-response genes preferential targets for ATF1 over cyclic-AMP response genes. The effect of this reciprocity is that each adaptation (weakly) suppresses the other. This may be important in advanced plaques with both non-resolving inflammation and IPH, as it means that IPH may worsen the deficiency in inflammation-resolution.

The role for histone acetylation is pharmacologically exciting as HDAC inhibitors are entering clinical practice^58^. The gene-activating histone signal H3K9Ac was found early on *HMOX1* in the hemin-response, and the broad-spectrum HDAC-inhibitor TSA increased both SMARCA4 recruitment to *HMOX1* and *HMOX1* mRNA expression. Following on from this, the HDAC3 inhibitor RGFP966 synergised with both hemin-induced *HMOX1* and PGI_2_-induced *NR4A2*. It will be important to equip macrophages to remove *both* IPH and inflammation, and likely also lipids, to better treat atherosclerosis. The beneficial effects of genetic *Hdac3*-deficiency *in vivo* in atherosclerosis in mouse models have been previously published^59^. This work suggests that new drugs targeting HDAC isoforms or similar epigenetic tags may show promise in re-instructing macrophages that have become specialised for lethality into ones that are dually equipped to clear arteries of hemorrhage and inflammatory cell debris.

### Summary

Data shown here point to a role for the SWI/SNF chromatin remodeller SMARCA4 in switching between expression and activity of effector enzymes *HMOX1* and *PLA2G7*, and between phagocytic clearance of oxidized RBCs and apoptotic WBCs. This likely reflects a reciprocal relationship between macrophage specialisation for inflammation-resolution and hematoma-resolution. It is mediated by chromatin remodelling prior to transcription factor recruitment, which alters the preferred binding sites for p-ATF1 from genes that coordinate leukocyte disposal and those that promote erythrocyte disposal. Since the remodeller is shown here to be independent as a risk gene for atherosclerosis and mediates functional switching between alternative macrophage pro-clearance phenotypes, these data form evidence of its pathophysiological importance in atherosclerosis. The reciprocity is likely to be overcome by HDAC3 inhibition, making this an attractive therapeutic approach.

### Conclusions

Several SNPs in the vicinity of *SMARCA4* have been associated with cardiovascular disease. Here, we demonstrate genetic independence of the *SMARCA4* SNP association from the *LDLR* SNP. We outline a mechanism in which metabolic-driven chromatin remodelling modulates preferred TF binding sites, so switching macrophage phenotype from inflammation-resolution to IPH-resolution. This mechanism is not well described in the literature. We show that the HDAC3 inhibitor RGFP966 may synergise with both responses. Further definition of this mechanism and potential translational outcomes with may eventually help continue to improve treatment of atherosclerotic cardiovascular disease by new and better-targeted anti-inflammatory drugs, possibly HDAC3 inhibitors.

## Financial Support

JJB was funded by a BHF Senior Clinical Research Fellowship FS13/12/30037, BHF Project Grants PG/15/57/31580, PG/17/71/33242, PG/21/10422). Some of the imaging was carried out with instruments in FILM facility, Imperial College London. The Facility for Imaging by Light Microscopy (FILM) at Imperial College London is part supported by funding from the Wellcome Trust (grant 104931/Z/14/Z) and Biotechnology and Biological Sciences Research Council (BBSRC) (grant BB/L015129/1). This work was supported by the NIHR Imperial Biomedical Research Centre (BRC). The views expressed are those of the author(s) and not necessarily those of the NIHR or the Department of Health and Social Care. SEH acknowledges funding from the BHF (PG08/008) and the UCL Biomedical Research Centre.

*In memoriam.* This manuscript is dedicated to the memory of Professor Justin Mason (co-author) (1961-2022) a supremely valued scientist, doctor, colleague, friend and mentor. Amongst many other things, he introduced Heme Oxygenase research to NHLI, so without him, this manuscript would never have existed

## Author contributions

JJB conceived the study, directed the study and carried out the bulk of the experiments. LC carried out the cyclic-AMP RNA-Seq experiment. DC, SS, AT, and FC carried out individual RT-qPCR experiments. SH carried out the genetics. KML, DOH and JCM provided intellectual input or critically commented on the manuscript.

