## Supplemental for "Hemin-driven chromatin remodelling by atherosclerotic risk gene *SMARCA4* switches human blood-derived macrophages from leukocyte disposal to erythrocyte disposal"

**Supplemental Text**

**Methods**

**Cell culture: macrophage culture and stimulation**

Cell culture was exactly as previously described[1]. Macrophage culture from humans and from mice was exactly according to our previous methods.

Human peripheral blood was obtained with informed consent and institutional ethical approval (VAS_JJB_17_076). PBMCs were prepared by density-gradient centrifugation and monocytes purified by adhesion as before. Human aseptically prepared LDL was purchased from Calbiochem (437644-10MG) and oxidatively modified with hypochlorite 1mM (from Sigma-Aldrich 425044-250mL) exactly as in our previous papers.

**Gene knockdown**

siRNA was carried out according to our previous methods. The si-RNA oligos were from Dharmacon and were pools of 4 sequences and were controlled for with non-targeting si-RNA from the same manufacturer at the same concentration and prepared in the same way at same time. All si-RNA were stored at -80°C. Si-RNA was made into complexes with Interferin (PolyPlus, Strasbourg, France, 409-10) by incubation of 100 pmol siRNA in 1μL with 1μL Interferin for 5-10 min. The complexes were then diluted in 100μL medium and added to a well of cells in 1 mL medium with 10% serum for 24 hours, at 100pmol siRNA per $10^5$ cells in each well. The si-RNA complexes were incubated with the macrophages for 18-24h before stimulation. Where multiple wells were treated, the incubation volumes of the mix were scaled by direct proportion.

**Bone marrow macrophage culture**

Mice were maintained on chow in a specific pathogen free facility in individually vented cages in Imperial College facilities in accordance with all institutional, national and

European guidelines. All experimental mice were of ages 4-6 months and of sexes 1:1 male: female. No statistically significant difference was observed with age or sex. Bones were collected aseptically from AMPK-KO or matched littermate control mice and flushed with PBS to collect bone marrow cells. First, legs were detached from the carcass, the femur was separated from tibia and fibula, and the ends of the femur were cut using a no. 20 size scalpel. Then, bone marrow was flushed out of each femur and tibia using a 25G needle on a 1ml syringe containing DMEM media (21909-035) with 10% FCS and 10% L929-conditioned media. The flushed bone marrow (in approximately 4ml media) was made up to 50ml of DMEM media + 10% FCS + 10% L929 media. The 50ml media containing bone marrow cells was divided into 4-6 T75 flasks and cultured at 37ºC, 5% $CO_2$ for 6 days in 10% FCS DMEM supplemented with antibiotics, L-Glu and 10% L929-conditioned medium (a source of M-CSF). Cells were then washed and adherent cells recovered by scraping into medium, counted and transferred to fresh 10% FCS DMEM in 24-well plates or 96-well plates.

**Treatments**

Hemin was from Sigma-Aldrich (Fluka 51280) and was aseptically prepared in DMSO (D2650) at 25mM and sterile-filtered and endotoxin-tested. Stock solutions in DMSO were stored at -80°C and thawed directly before use. All stock solutions were diluted >1000-fold when adding to cultures.

SB747651A was purchased (Tocris #4630) and made up to 25mg/mL in PBS (60mmol.$L^{-1}$), aliquoted and stored at -80ºC. Aliquots were diluted to 1mmol.$L^{-1}$ in medium and then at 1:100 (10µL in 1mL) to a final culture concentration of 10µmol.$L^{-1}$.

dbc-AMP was purchased (Merck-Millipore 28745-100MG) and made up to 0.1M in tissue culture water (Sigma-Aldrich W4502-1L) and diluted 1:100 to 100µmol.$L^{-1}$ final concentration in culture medium.

Trichostatin-A (TSA) was T8552 from Sigma-Aldrich made up to 1mmol.$L^{-1}$ in DMSO (Hybridoma Grade, Sigma Aldrich D2650), diluted to 100nM final concentration in medium (as 1µL to 100µL in IMDM, then 10µL to 1mL final).

Apremilast (Enzo #BV-2352-5) was purchased and made up to 80mg/mL stock in tissue culture water (Sigma-Aldrich W4502-1L) and then diluted to final concentration 100nmol.$L^{-1}$ in culture medium.

**RT-qPCR**

Macrophages were lyzed in a guanidinium-based buffer (RLT buffer, RNA-Easy Mini kit, 74104, Qiagen Manchester, UK) and RNA purified by silica-resin affinity (according to manufacturer's instructions). Reverse transcription was with Invitrogen Superscript-II, following manufacturer's instructions (Invitrogen, Life Sciences, Paisley, UK, 18064022). Quantitative polymerase chain reaction (qPCR) was with ausing a BioRad iCycler CFX96 real time PCR, MesaGreen mastermix (Sybr Green/Taq, Eurogentec, Southampton, UK, RT-SY2X-03+WOUFL) and 100fmol.µ$L^{-1}$ custom primers (synthesized by MWG Biotechnology, Ebersberg, Germany). Primers were designed by a Primer3 algorithm accessed via the NCBI website (www.ncbi.nlm.gov). Additional validation was on a further five donors over a longer time course, using qPCR by the same methods. Primer sequences are in the Supplemental Table.

**QPCR**

Other than exceptions below, most qPCR was carried out on a BioRad CFX96 instrument. Stock primers and cDNA were diluted by 1:10 (5ul primers/cDNA added to 45ul molecular biology water (Sigma-Aldrich W4502-1L)). Master-mix was made up for the appropriate number of wells, with containing the following:

| | |
|---|---|
| MESA Green | 7.5ul |
| Forward Primer | 0.25ul |
| Reverse Primer | 0.25ul |
| Water | 4.0ul |

Total 12ul per well

2ul of diluted cDNA added to each well. Each sample was run in triplicate.

The qPCR protocol was as follows:

1. 95ºC for 3 mins
2. 95ºC for 10 secs
3. 60ºC for 1 min
4. Repeat (Steps 2-3) 40 times
5. Plate read melt curve 56ºC to 95ºC for 5 secs

For primers producing a product longer than 150bp, the following protocol was used:

1. 95ºC for 3 mins
2. 95ºC for 10 secs
3. 60ºC for 2 mins
4. Repeat (Steps 2-3) 40 times
5. Plate read melt curve 56ºC to 95ºC for 5 secs

Mesa Green is a SyBr Green mastermix with ROX, (Eurogentec, RT-SY2X-03+WOUFL). Primers were custom-ordered from MWG Eurofins (Supplemental Table). For ChIP reactions, qPCR was carried out for 60 cycles.

The qPCR reactions in Figure 6B-E and Figure 5A were carried out using a QuantStudio 6 (ThermoFisher) and Luna qPCR mix (M3003L, NEB) using the following settings:

384-well plate

4µL Luna per well

4µL sample per well

1. 1.6ºC/s
2. 50ºC 02:00
3. 1.6ºC/s
4. 95ºC 08:00
5. 1.6ºC/s
6. 95ºC 00:15
7. 1.6ºC/s
8. 60ºC 01:00
9. Camera
10. FOR N=60 cycles, GOTO 6
11. 1.6ºC/s
12. 95ºC 00:15
13. 1.6ºC/s
14. 60ºC 01:00
15. 0.05ºC/s (Camera)
16. 95ºC 00:15

**Chromatin immunoprecipitation - qPCR**

Macrophages were washed once with 500µL / well PBS at 4ºC, crosslinked in 1% Formaldehyde Molecular Biology Grade (from Sigma-Aldrich F8775-25mL diluted in PBS (20012-019 Gibco ThermoFisher)) for 15mins at 4ºC; washed once in 500µL / well PBS; then crosslinking stopped with 0.1M Glycine for 15mins at 4ºC; then aspirated to dryness and stored at -80ºC until further analysis.

Then the wells were thawed and lysed for 20 minutes in the presence of 100µL/well nuclear lysis buffer AM-1 (Active Motif #100566) supplemented with Protease Inhibitor Cocktail (Active Motif #100546) and Phosphatase Inhibitors (Active Motif #102146) and then scarped into 100µL of the lysis buffer. The lysate was passed x10 through a 27G insulin syringe (Becton Dickinson) and sonicated in a Covaris Sonicator (Cavaris S220 Ultra Sonicator) at settings Peak Power 105, Duty Factor 2.0, Cycles/Burst 200, Duration 600s.

The sonicated lysates were then diluted in ChIP buffer of composition: 20mM Tris pH7.2 (1mL of 1.0M Tris pH7.2 (from Sigma-Aldrich 41573-1L-F) in 50mL Molecular Biology water (Sigma-Aldrich W4502-1L)), 150mM NaCl (3mLs of 2.5M NaCl (Sigma-Aldrich S3104-500G 2.5M in Molecular Biology water (W4502-1L)) in 50mL Molecular Biology water (Sigma-Aldrich W4502-1L)). Antibody was added (purified rabbit monoclonals against respective targets, 0.2µg.mL$^{-1}$ final, (Abcam), or Abcam control monoclonal rabbit IgG to the same concentration) and rotated at 4ºC for 16h in 1.5mL eppendorf-type microcentrifuge tubes (VWR or Sarstedt). Then 4µL per eppendorf of 1:10 diluted Protein-A magnetic beads (ThermoFisher Invitrogen 10001D) in Molecular Biology grade water were added and rotated for 2h at room temperature (15ºC). the beads were separated with a Neodymium magnet, and transferred to

100µL of Proteinase K (PK) (Proteinase K (Promega #P81025) in PK buffer (SDS 0.1% (Molecular Biology Grade, from Sigma-Aldrich 05030-500mL-F), 10mM Tris pH8, $CaCl_2$ 1mM (from Sigma-Aldrich BioUltra Molecular Biology Grade, 21115-100ML). The PK digestion was at 65ºC for 18h. Then PK reactions were extracted with 100µL Phenol:Chloroform:isoamyl alcohol (Sigma-Aldrich 77617-100mL) the aqueous layer removed and precipitated with 1:20 volume of sodium acetate 3.0M (Molecular Biology Grade, Ambion AM9740) and 2x volume of Molecular Biology Ethanol (Fisher BP2818-500). After 48h at -80ºC, the DNA was spun down at 18,000G for 3h and ethanol supernatant removed and pellets evaporated to dryness in a laminar flow hood. The DNA pellet was then resuspended in 400µL Molecular Biology water (Sigma-Aldrich W4502-1L).

**Chromatin conformation capture analysis - qPCR**

Macrophages were washed once with 500µL / well PBS at 4ºC, crosslinked in 1% Formaldehyde Molecular Biology Grade (from Sigma-Aldrich F8775-25mL diluted in PBS (20012-019 Gibco ThermoFisher)) for 15mins at 4ºC; washed once in 500µL / well PBS; then crosslinking stopped with 0.1M Glycine for 15mins at 4ºC; then aspirated to dryness and stored at -80ºC until further analysis.

Macrophages were then thawed in 10µL buffer AM-1 with added 0.3% SDS (Molecular Biology Grade, from Sigma-Aldrich 05030-500mL-F). They were lysed at in a shaker at 900rpm 37ºC for 2h. Then, the lysates were diluted and incubated in Bgl2 (NEB) in 1:10 of 10x Bgl2 buffer (NEB) 50µL and 1.2µL per reaction of 50,000u/mL Bgl2, on a ratio of $10^5$cells/4u Bgl2, for 24h. Then 4µL 20% SDS (Molecular Biology Grade, from Sigma-Aldrich 05030-500mL-F) were added to the 50µL reaction volume to inactivate the Bgl2 and the mix was diluted to 700µL in T4 ligase buffer.

T4 Ligase (Promega M1808) 3u/µL 7L/reaction was added in T4 ligase buffer and incubated at 16ºC for 36h. Next, the reaction mix was extracted in an equal volume of Phenol:Chloroform:isoamyl alcohol (Sigma-Aldrich 77617-100mL), the aqueous layer removed and precipitated with 1:5 v/v 3.0M sodium acetate (Molecular Biology Grade, Ambion AM9740) and 5:1 v/v ethanol (Molecular Biology Grade (Fisher BP2818-500)). The DNA was precipitated at -80ºC for 48h, spun for 3h at 18,000G, and supernatants removed. The pellets were dried in a laminar flow hood for 48hand then dissolved in Molecular Biology water (Sigma-Aldrich W4502-1L).

**RNA-Sequencing and analysis**

**Reagents**

dbcAMP (Merck-Millipore 28745-100MG); Monarch total RNA miniprep kit (T2010S, New England Biolabs, Hitchin, UK); RNase-free water (Thermo Scientific, Loughborough, UK); RNaseZAP (Thermo Scientific, Loughborough, UK); Agilent RNA 6000 Nano Kit, containing RNA dye concentrate, RNA 6000 nano gel matrix, RNA 6000 nano ladder, RNA 6000 nano marker (Agilent, Santa Clara, California, USA); Ampure XP beads (Beckman Coulter, Brea, California, United States); EB buffer (Qiagen, Manchester, UK); Luna qPCR Reagent (M3003L NEB)

**Sample preparation and selection**

HMDMs isolated from 4 healthy donors were subjected to 3 treatments *in vitro,* 100µM dbcAMP or vehicle (water (Sigma-Aldrich W4502-1) for 1 hour. Cells were plated in 24-well plates at a density of 0.75x$10^6$ cells/well with 12 wells per treatment. A separate set of wells were prepared to analyse HMDM purity by measuring CD14 and CD16 expression via flow cytometry as described in (reference section). After stimulation, media was removed and cells were lysed with 300µl RNA lysis buffer per treatment, followed by scraping with a rubber policeman. Total RNA was extracted using Monarch total RNA miniprep kit, as described in (reference section). Isolated total RNA was eluted into 40µl RNase-free water and then divided between 3 x 1.5ml tubes as follows, 30µl for RNA sequencing, 5µl for RNA quality control testing and 5µl for RT-qPCR. Samples were then immediately placed in -80$^{o}$C for storage. 5µl of RNA assigned for RT-qPCR was processed as described in (reference section) and relative NR4A2 mRNA levels measured. Samples were rejected from the study if less than 75% of cells expressed CD14 or NR4A2 mRNA induction in response to dbcGMP was below 1.5-fold.

**RNA quality control**

5µl aliquot of total RNA assigned for quality control was analysed using Agilent RNA 6000 Nano Kit on a 21000 bioanalyzer (Agilent, Santa Clara, California, USA), following manufacturer's instructions. To decontaminate the bioanalyzer electrode, one well of the electrode cleaner chip was filled with 350µl RNaseZAP and placed into the bioanalysed for 1min with the lid closed. A second cleaner chip filled with 350µl RNase-free water was placed in the bioanalyzer for 10sec. The cleaner chip was removed and the lid of the bioanalyzer was then left open for 10sec to allow the water to evaporate. Filtered gel matrix was prepared by centrifuging 550µl RNA nano 6000 gel matrix in a spin filter column at 15000g for 10mins and then divided into 32µl aliquots. To prepare the gel-dye mix, RNA dye concentrate was vortexed for 10sec and spun down briefly. 0.5µl of RNA dye concentrate was added to a 32µl aliquot of filtered gel, vortexed thoroughly and then centrifuged at 13000g for 10min at room temperature. RNA samples and RNA 6000 nano ladder were denatured by incubation at 70$^{o}$C for 2mins and then placed on ice. The nano chip was primed with 9µl of gel-dye mix and 30sec of applied pressure using a chip priming station (Agilent, Santa

Clara, California, USA). A further 9µl of gel-dye mix was applied to the 2-remaining gel-dye wells, 5µl of RNA 6000 nano marker was applied to all remaining wells and 1µl of RNA 6000 Ladder was applied to the ladder well. 1µl of each RNA sample was added to its assigned sample well and then the nano chip was vortexed for 1min. The nano chip was placed in the bioanalyzer and analysed using the Eukaryote Total RNA Nano Series II setting. A set of samples from a single donor were rejected if one sample produced a RIN score of <7.0, an RNA quantity of <200ng or an RNA concentration of <10ng/µl.

**Library preparation and RNA-sequencing**

Library preparation and RNA sequencing was performed externally using the BGISEQ-500 platform (BGI Genomics, Tai Po, Hong Kong). 30µl aliquots of total RNA samples assigned for RNA sequencing were transferred from -80$^{o}$C storage directly to dry ice and sent to BGI Genomics by aircraft. BGI Genomics performed an independent quality control of the RNA samples, matching the bioanalyzer results. The following is a summary of the key steps in the BGI Genomics library preparation and sequencing protocol. mRNA was purified from total RNA samples using oligo(dT)-attached magnetic beads. mRNA molecules were then enzymatically fragmented into short strands. First-strand cDNA was generated using random hexamer-primed reverse transcription, followed by second-strand cDNA synthesis. The synthesised cDNA was subjected to end-repair, followed by 3’ adenylation. Adapters were ligated the ends of the 3’ adenylated cDNA fragments and amplified via PCR. PCR products were purified using Ampure XP beads and dissolved in EB buffer. Library quality was validated using a 2100 bioanalyzer (Agilent, Santa Clara, California, USA). The double stranded PCR products were heat denatured and circularised using a splint oligo sequence. The single stranded circularized cDNA library was amplified using phi29

DNA polymerase producing DNA nanoballs (DNB) which had more than 300 copies per single-stranded circularized cDNA molecule. DNBs were loaded onto a patterned nanoarray and RNA-sequencing carried out using the BGISEQ-500 sequencer. Using a sequencing by synthesis method, pair-end 100 base pair reads were produced with 30 - 35 million reads per sample. BGI provided adapter-trimmed raw read files only and did not perform any further bioinformatic data processing or analysis.

**Bioinformatic analysis of large genesets**

**Software**

PARTEK (https://www.partek.com/); PANTHER-GO (http://pantherdb.org/); GSEA (http://www.gsea-msigdb.org/gsea/login.jsp); InnateDB (https://www.innatedb.com/); STRING (https://string-db.org/); PASTAA (http://trap.molgen.mpg.de/PASTAA.htm); oPOSSUM (http://opossum.cisreg.ca/); Venn Diagram Generator (http://www.pangloss.com/seidel/Protocols/venn.cgi); X2K web (https://amp.pharm.mssm.edu/X2K/); ChEA3 (https://amp.pharm.mssm.edu/chea3/).

**Analysis**

RNA sequencing read files were quality checked, processed and analysed using PARTEK software. BGI provided Fastq files that contained adapter-trimmed 100 base pair reads. After read quality was tested, alignment was performed using the STAR aligner tool. The aligned reads were quantified to the PARTEK E/M annotation model producing the number of counts per gene. Gene counts were normalised to read depth per sample, using the counts per million (CPM) method. Due to high variation between human donors, donor variability was removed using the remove batch effect tool. The statistical comparison test, gene-specific analysis, was performed between samples resulting in a fold change value, p-value and FDR-value per gene per comparison.

Downstream analyses were performed on a series of web-based software: heatmaps was produced using PARTEK, pathway and gene ontology analyses were investigated using PARTEK, DAVID and InnateDB, transcription factor binding site enrichment was tested using X2K web, oPOSSUM and ChEA3 and transcription factor networks were explored using STRING.

After RNA-sequencing was performed, read files were quality checked, processed and analysed using PARTEK software. Read quality tests showed that that both the average base quality score per position and the average quality score per read was above 30. Base quality score is a measure of the probability that the correct base has been called and a score of 30 translates to a 99.9% base call accuracy. This indicates read files were of a good quality. Alignment was performed using the STAR aligner tool. Post-alignment quality tests demonstrated that at least 75% of reads were aligned to the genome per sample and that samples had an average read depth of a covered region of at least 30. The aligned reads were quantified to the PARTEK E/M annotation model producing number of counts per gene. Gene counts were normalised to read depth per sample, using the counts per million (CPM) method. The statistical comparison test, gene-specific analysis, was performed between the 2 test groups resulting in a fold change value, p-value and FDR-value per gene.

**Statistics**

Data were graphed in stem-and-leaf plots and examined visually for distribution, and tested for normality using Shapiro-Wilk when $n \geq 5$. SigmaPlot was used for routine analysis. SigmaPlot reports exact p-values down to $p<0.001$. GraphPad Prism was used for exact p-values down to $p<0.0001$. SPSS was used for remaining exact p-values. SPSS was used for time-series analysis.

Animals were randomized using the randomization function in Microsoft Excel (=RAND()). Alternate allocations were made for values over or under 0.5.

Data that passed normality by Shapiro-Wilk, visual inspection, general acceptance of normal distribution, and n≥5, were tested as follows. Inherently paired data were tested using Student's paired t-test. Non-paired data in two groups were tested using Student's t-test. Data in several groups, were tested using One Way ANOVA and post-tested using Holm-Sidak correction for multiple simultaneous comparisons. Time-series data were significance tested using Repeated Measures One Way ANOVA.

Data that failed normality testing were significance tested as follows. Data in multiple groups were tested using One Way ANOVA on Ranks (Kruskal-Wallis), with Dunn's post-test. Unpaired data in 2 groups were tested using Wilcoxon Rank Sum Test (Wilcoxon-Mann-Whitney).

No corrections for multiple testing were made across tests (*i.e.* only within-test corrections were made), When any comparison is not shown it was not run. In several figures, only 1 comparison between 2 treatments was key (*e.g.* Day 9, KO vs control) and this comparison was chosen in advance. Whenever a multiple comparison procedure is referenced, a post-hoc p-value correction was used

**Network analysis**

Gene expression data were input to STRING (https://string-db.org/). The settings were left to default other than the use of one round of 'more'. The network was then exported as an adjacency matrix and imported into Cytoscape for visualisation (https://cytoscape.org/).

**ELISA**

ELISAs for $LTB_4$, $RvD_1$ and $PGI_2$ were carried out on saved culture supernatants. Cayman Chemicals 96-well colorimetric ELISA kits were used, exactly according to manufacturer’s instructions (CAY520111, CAY500380, CAY50110). The same kit was used for both species (human and murine eicosanoids are chemically identical). Absorbance was measured at the manufacturer’s recommended wavelength, using a Tecan SpectraFluor 96-well plate reader. ELISAs for serum Amyloid A (SAA) were purchased as the DuoSet kit for mouse SAA (R&D Systems DY2948-05) and were used as per manufacturer’s instructions, including the calibration in duplicate and samples in technical triplicate.

**Western blotting**

RAW cells were lysed in Laemmli buffer (10%SDS (Sigma-Aldrich 71725), 20% glycerol (Sigma-Aldrich G5516, TrisHCl-pH6.8, 1:100 phosphatase inhibitors (Active Motif phosphatase inhibitor cocktail 37493) no bromophenol blue or 2-ME)) and subjected to SDS-PAGE with a Novex system (Invitrogen kit NP030A)(Invitrogen NuPAGE precast BisTris 4-12% gradient (NP0321BOX), MOPS running buffer NP0001) and Rainbow Markers RPN800E, electrotransferred using Novex to Immobilon-FL (ThermoFisher Pierce 88518). Membranes were blocked for 10mins with 10% BSA TBS and incubated overnight with 1:1000 anti-HO-1 (rabbit monoclonal Abcam Ab52947) or anti-beta-actin (Abcam Ab6276) and then developed respectively with anti-rabbit or anti-mouse Peroxidase-labelled (Dako P0447 and P0448) and ECL-Plus (ThermoFisher Pierce 32132) and Amersham Hyperfilm (VWR 28-9068-37) and exposed over a log range, and the exposure with highest signal:noise ratio used.

**Functional analyses**

**PAF-AH assay**

hMDM were cultured as above and lysed in Molecular Biology grade water (200µL) per well and stored at -80°C. lysates, 100µL was used per assay, by minor modification of published methods[2,3,4]. NPS-PC (Merck 810857P) was added at 10µM and absorbance measured at 400nm.

**HO-1 assay**

hMDM were cultured at confluence (2x$10^4$ cells per well) in a 96-well plate and loaded with hemin by incubation with 20µM hemin 60 mins, then absorbance was measured over a spectrum every 5nm from 350-600nm every 24h. The rate of change of decrease of the hemin peak was calculated and the rate of change of hemin in cells calculated by reference to hemin standards.

**Phagocytosis**

hMDM were cultured in 8-well chamber slides at $2x10^4$ cells per well, with si-RNA for 48h, then $PGI_2$ (100nM) or hemin (10µM) were added for a further 48h. Then fluorescently-labelled oxidatively damaged erythrocytes or apoptotic leukocytes were added, at 10:1 targets:macrophages i.e. $2x10^5$ cells per well. Coincubation was over 16h, then the supernatants were gently resuspended, and collected. The cell monolayer was then fixed and counterstained with DAPI.

RBCs were oxidatively damaged by dilution 1:5 in PBS to 5mL; then 2 additions of 50µL $H_2O_2$; incubated for 30mins 37°C, centrifuged 800G 10mins; then 100µL of pellet were diluted 1:10 in PBS, 1:100 1mg/mL 1’,1’-dioctadecyl-3’,3’,3’,3’-tetramethylindocarbocyanine perchlorate (Sigma-Aldrich 42364, di-I, a green fluorescent membrane-intercalating dye), and then washed once by centrifugation and resuspension.

Apoptotic leukocytes were from the RAW264.7 cell line, grown in continuous culture in 10%FCS IMDM, grown to confluence in a T75 flask, scraped and transferred to PBS, washed in PBS by 3 rounds of centrifugation and resuspension, then irradiated under a UV lamp in PBS for 4h, and collected and di-I-stained as for the RBCs. These conditions produce predominantly post-apoptotic secondary necrosis

**Supplemental Figure Legends**

**Supplemental Figure 1**

Expression level of SMARCA4 and LDLR in different tissues. Data from GTex portal

**Supplemental Figure 2 Expression of key genes in response to cyclic-AMP or hemin by RT-qPCR.** X-axes, time in hours after hemin or cAMP treatment up to 96h (4 days) (log scale). Y-axes, induction of indicated genes in fold from baseline, calculated by $2^{-\Delta\Delta Ct}$ method as standard. A, C, E, G – stimulation with 100μM dbc-AMP (n=8 donors). B, D, F, H – stimulation with 10μM hemin (n=8 donors). A, B – *NR4A2*, C,D – *FOS*, E,F – *HMOX1*, G,H – *PLA2G7*.

**Supplemental Figure 3 Expression of key genes in response to $PGI_2$ by RT-qPCR.**

X-axes, time in hours after 1μM $PGI_2$ (Iloprost) treatment up to 8h. Y-axes, induction of indicated genes in fold from baseline, calculated by $2^{-\Delta\Delta Ct}$ method as standard. A - *NR4A2; B – FOS,* C - *HMOX1*. Data are mean S.E. (n=6 donors).

**Supplemental Figure 4 Network analysis**

Genes were input to a protein-protein interaction database to establish relationships (https://string-db.org/). A,B, network diagrams for hemin (red) (A) and cyclic-AMP (blue) (B). Each point represents on gene plotted onto the STRING database and each line represents a known interaction between each gene and each other gene in the gene expression profile. A small number of nodes are highly connected: this is systematically graphed in (C). I, Y-axis, number of nodes (connected genes) in network; X-axis number of connexions, grouped on a logarithmic scale.

Superimposed are the genes that in the top two groups for connectedness for each of hemin and cyclic-AMP.

C, Y-axis, number of nodes with connection; X-axis, number on connections per node, binned into exponentially increasing ranges. For cyclic-AMP, there was a typical power-law relationship, that is, log node degree (number of connexions) versus number of nodes forms a straight line. With hemin, this relationship was atypical, with a bell-shaped graph, for unknown reasons. The network analysis allowed identification of the most highly connected genes, which are likeliest to be important regulators, shown superimposed. For dbc-AMP, these were TFs and endocrine/paracrine mediators: *FOS*, *MYC*, *TNF*, *VEGF*, *CXCR4*, *ATF3*, *DUSP1*, *CCR5*, *SOCS3*, *EGR1*, *SMAD3*, *CXCL2*, *HSPA8*, *FOSB*, *NR4A2*. With hemin, the highly connected nodes were predominantly chromatin-regulators: *CREBBP*, *POL2RL2*, *SNRPD2*, *EZH2*, *CCT2*, *SKP1*, *NOP58*, *HSP90*, *NPM1*, *POL2RJ*, *CCT6A*, *CCT4*, *H2AFV*.

**Supplemental Figure 5**

**A, diagram showing locations of primer pairs** (red) relative to *HMOX1* gene. Key loci and distances given as shown. Arrow, transcriptional start (approximate) and direction. Red oligo icons, 1-6, locations of oligos used in qPCR. Not to scale.

**B, 3C by qPCR,** using qPCR primers on cut and re-ligated gDNA to quantify proximity of the *HMOX1-4100* site to the TSS. Y-axis, fold change from baseline. X-axis, time-point (hours after hemin-treatment). *p<0.05, n=5 donors, ANOVA

**C, ChIP-qPCR for SMARCA4 occupancy** on *HMOX1*-4100. Y-axis, fold relative to baseline. X-axis, time-point (hours after hemin-treatment). *p<0.05, n=5 donors, ANOVA.

**D, ChIP-qPCR for pATF1 occupancy** on *HMOX1-4100*. Y-axis, fold relative to baseline. X-axis, time-point (hours after hemin-treatment). *p<0.05, n=5 donors, ANOVA

**Supplemental Figure 6**

Effects of prostacyclin ($PGI_2$) and specialised pro-resolving mediator Resolvin D1 ($RvD_1$) on cyclic-AMP responsive genes *FOS* and *NR4A2* in hMDM after 4h stimulation after 1 day in culture. Apm - apremilast (an inhibitor of cyclic-AMP phosphodiesterase (PDE4)), 1µM.  y-axes, fold induction from baseline. Apremilast pre-incubated for 30 mins. *, p<0.05, Repeated measures ANOVA.

A, $PGI_2$ 100nM, left graph *NR4A2*, right graph *FOS*.

B, $RvD_1$ 10nM, *NR4A2*, right graph *FOS*.

### Supplemental Figure 1

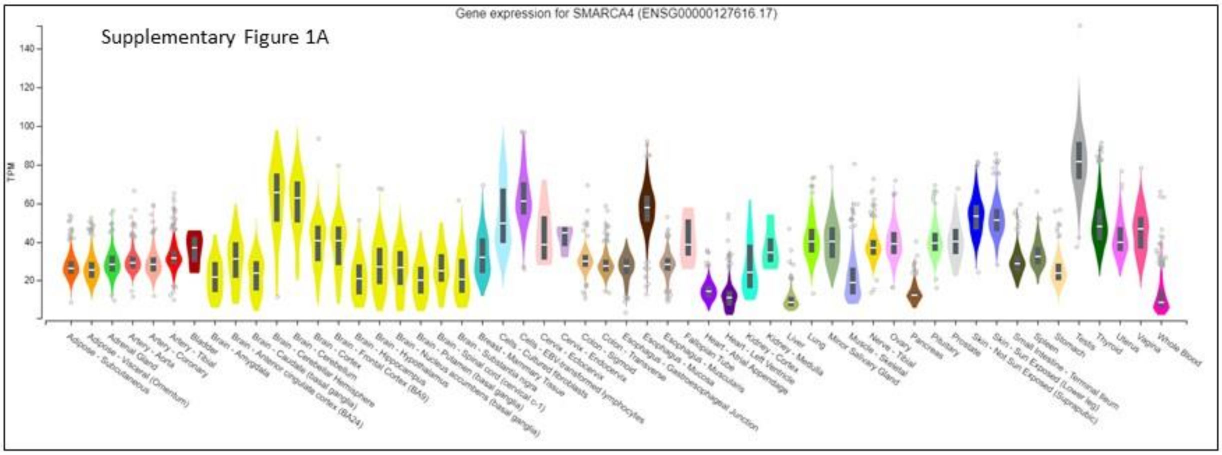


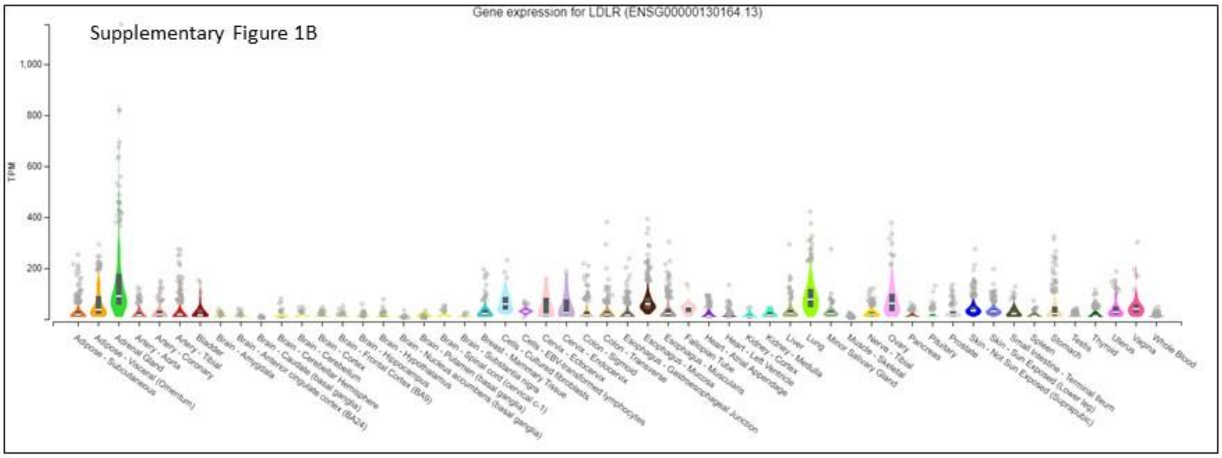


**Supplementary Figure 1**
Expression level of SMARCA4 and LDLR in different tissues. Data from GTex portal

### Supplemental Figure 2

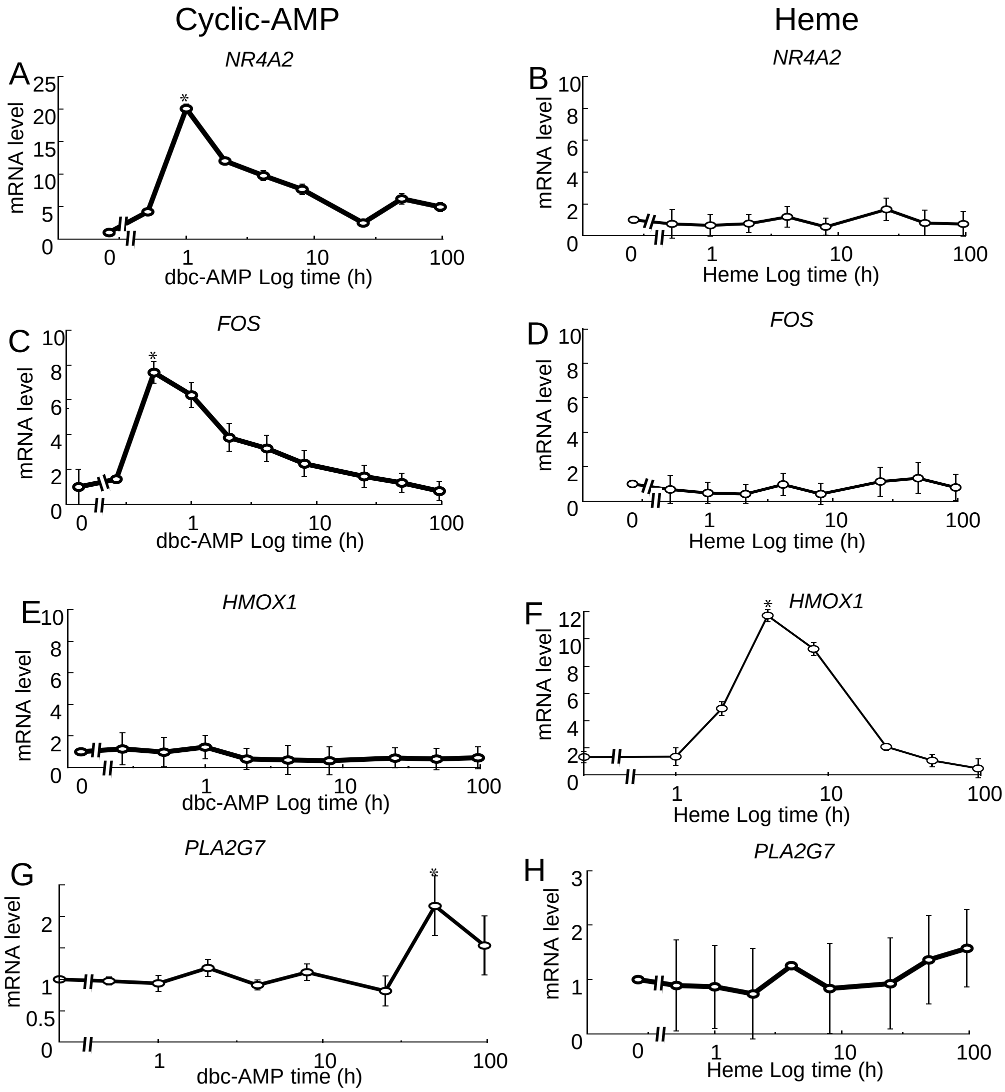

### Supplemental Figure 3

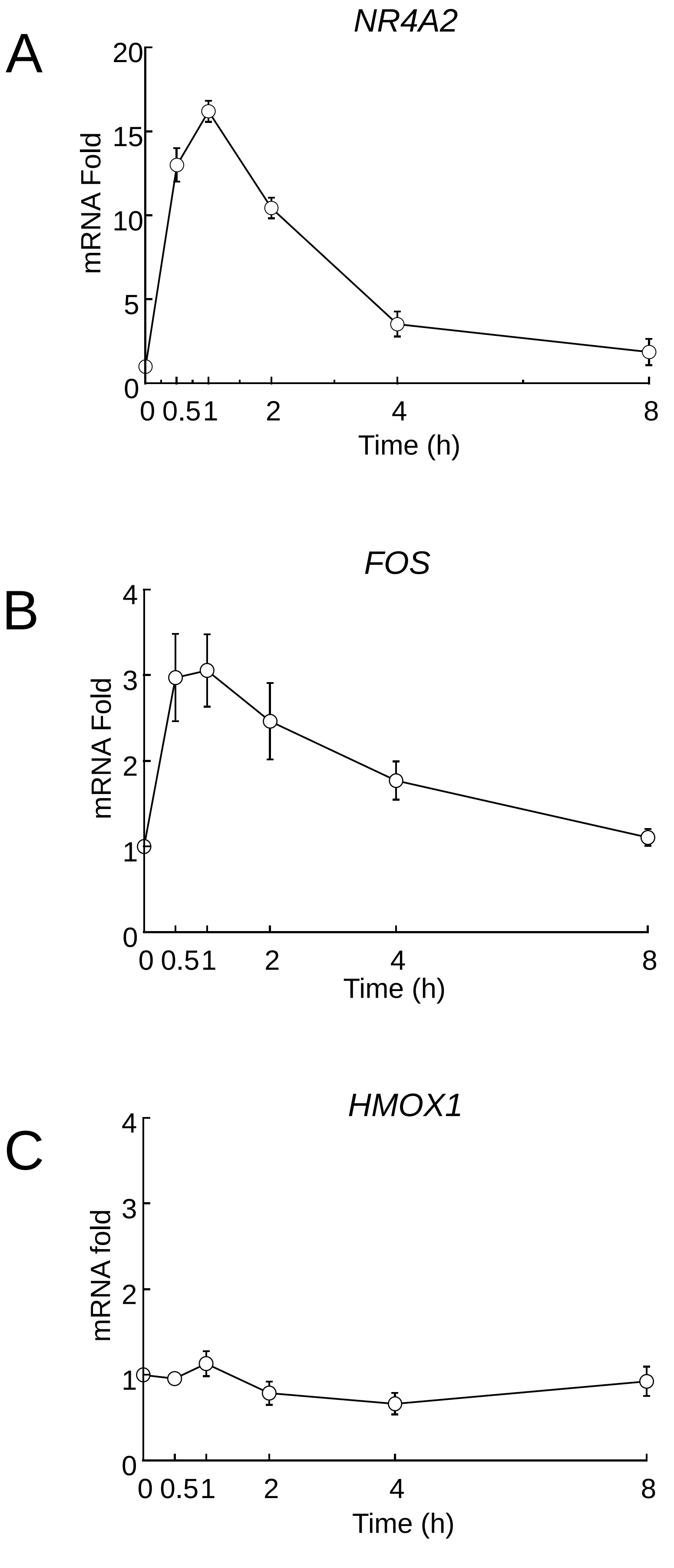

### Supplemental Figure 4

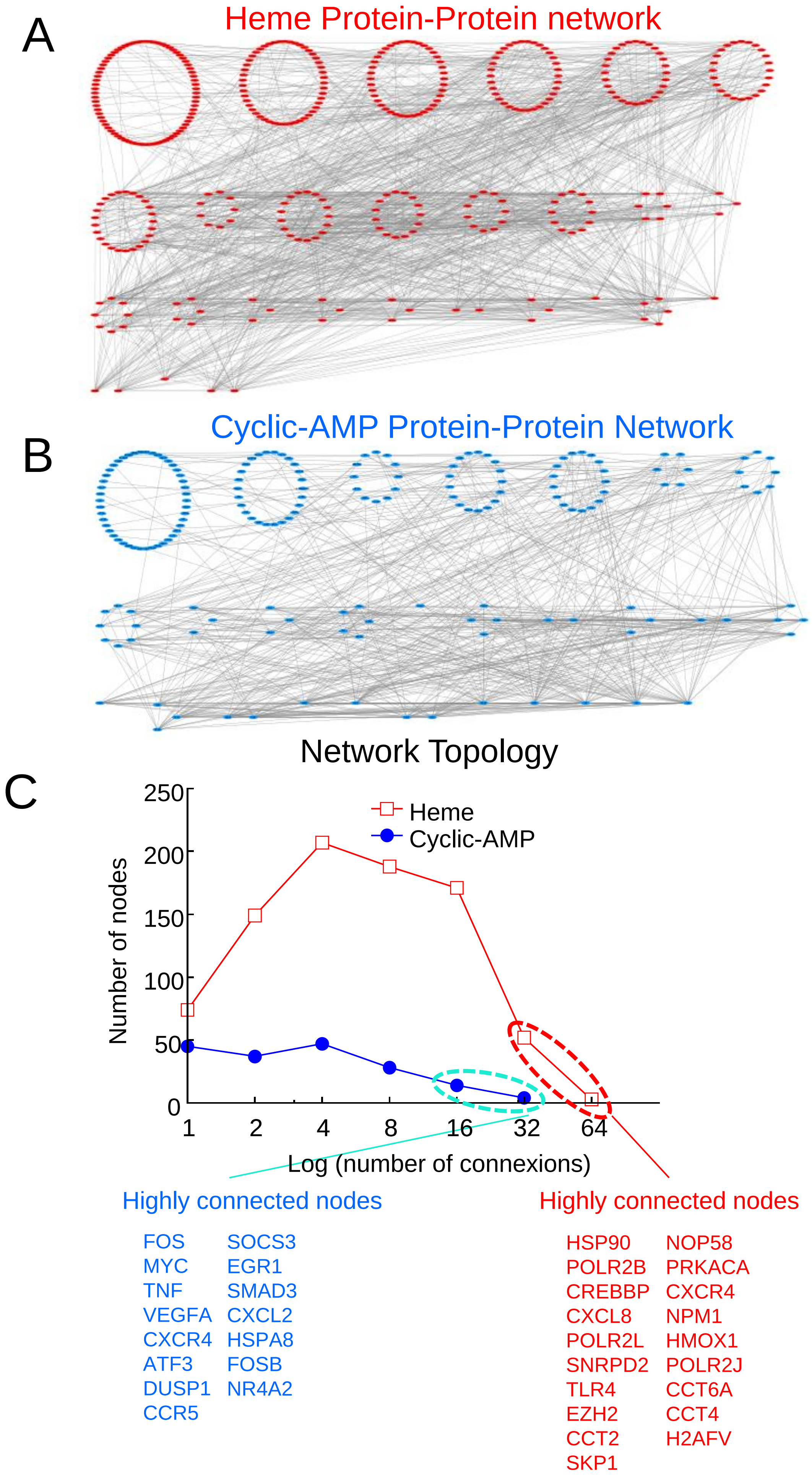

### Supplemental Figure 5

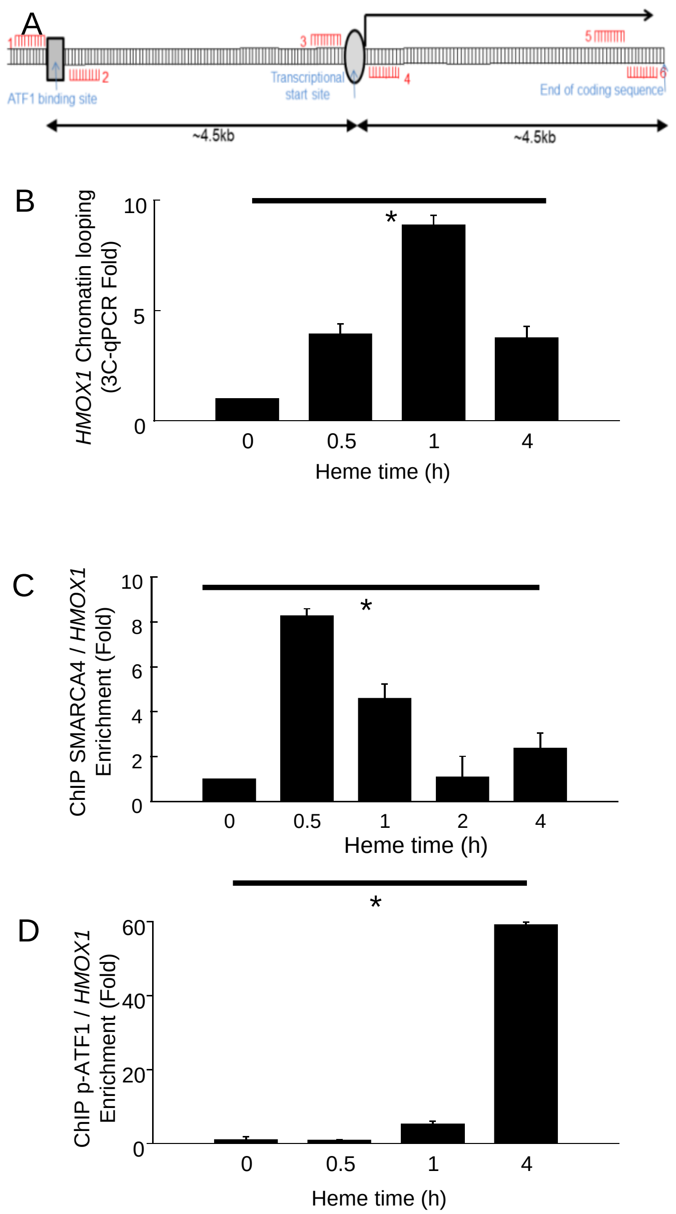

### Supplemental Figure 6

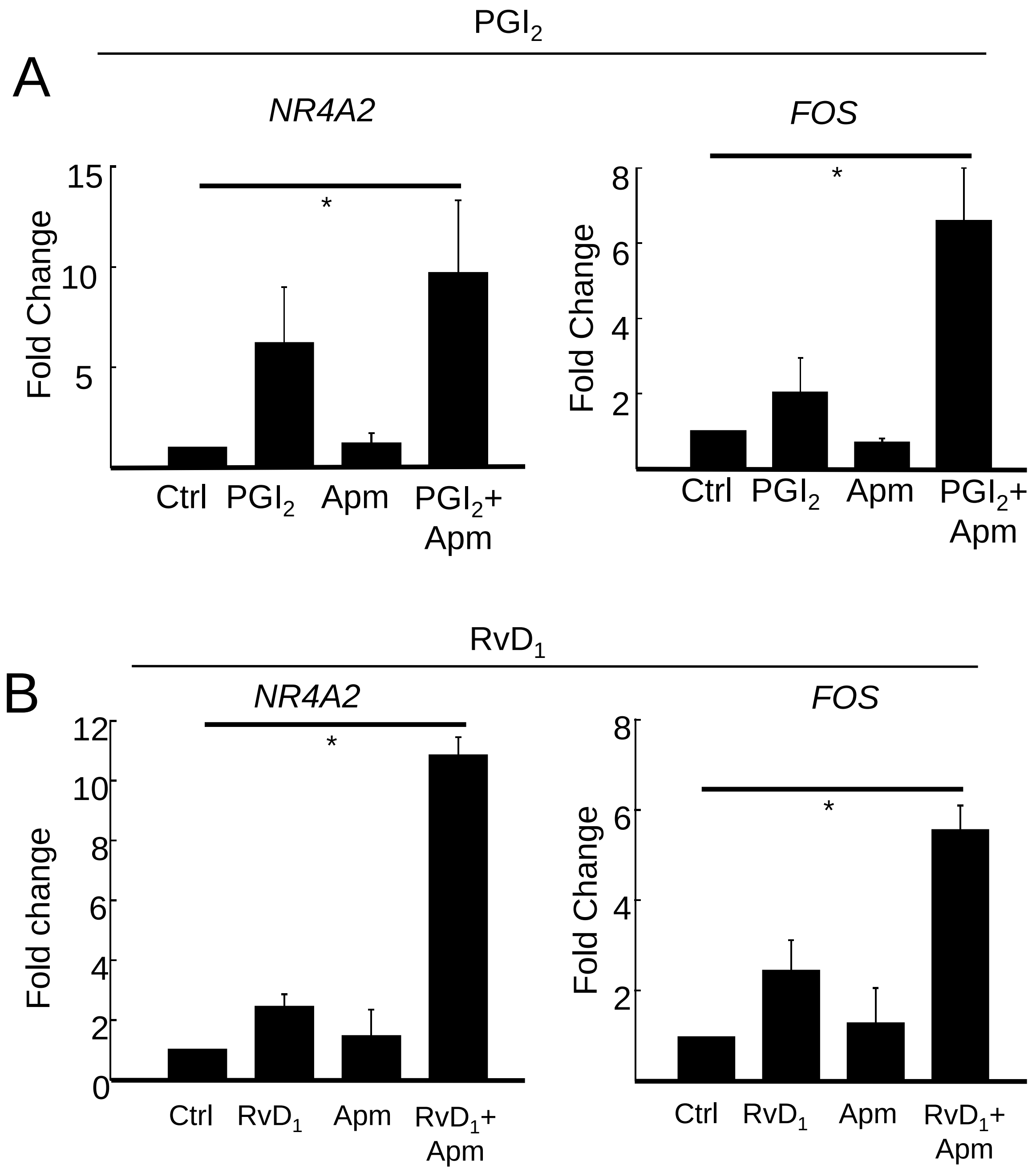
